# Native Electrospray-based Metabolomics Enables the Detection of Metal-binding Compounds

**DOI:** 10.1101/824888

**Authors:** Allegra Aron, Daniel Petras, Robin Schmid, Julia M. Gauglitz, Isabell Büttel, Luis Antelo, Hui Zhi, Christina C. Saak, Kien P. Malarney, Eckhard Thines, Rachel J. Dutton, Manuela Raffatellu, Pieter C. Dorrestein

## Abstract

Metals are essential for the molecular machineries of life, and microbes have evolved a variety of small molecules to acquire, compete for, and utilize metals. Systematic methods for the discovery of metal-small molecule complexes from biological samples are limited. Here we describe a two-step native electrospray ionization mass spectrometry method, in which double-barrel post-column metal-infusion and pH adjustment is combined with ion identity molecular networking, a rule-based informatics workflow. This method can be used to identify metal-binding compounds in complex samples based on defined mass (*m/z*) offsets of ion features with the same chromatographic profiles. As this native metal metabolomics approach can be easily implemented on any liquid chromatography-based mass spectrometry system, this method has the potential to become a key strategy for elucidating and understanding the role of metal-binding molecules in biology.

## Main

Life, as we know it, cannot exist without metals. Metals are essential cofactors in important biochemical reactions such as DNA replication and repair (iron and manganese), respiration (iron and copper), photosynthesis (iron and manganese), and biosynthesis of countless primary and secondary metabolites (iron, zinc, vanadium, molybdenum, magnesium, calcium, etc.). One common strategy for metal acquisition in microorganisms is through the production and secretion of small molecule ionophores that bind and form non-covalent metal complexes. These complexes bind to specific ionophore receptors for their uptake and release of metal into the cytoplasm. Siderophores are high affinity chelators of ferric iron (Fe^3+^) and include ferrioxamines (i.e. desferrioxamines B and E)^2–5^, catecholates (i.e. enterobactin)^2–4^, and carboxylates (i.e. rhizoferrin and aerobactin)^2–5^, while chalkophores such as methanobactins and SF2768 (a diisonitrile natural product) play the analogous role in binding cuprous copper (Cu^+^)^6,7^, and a number of zincophores have also been recently elucidated^8–11^. In addition to small molecule ionophores that play a role in microbial metal transport and homeostasis, metals can also serve as cofactors essential for the function of other small molecules. Examples include iron in heme and magnesium in chlorophyll, calcium and magnesium in metal-dependent antibiotics^12–14^, and cobalt in the vitamin cobalamin (vitamin B12).

Although metal-binding small molecules have a variety of biological functions and many potential biomedical applications, it is currently challenging to find metal-binding compounds present in complex mixtures such as microbial culture extracts^15^, dissolved organic matter^16^, or fecal extracts^17^, and it is additionally challenging to assess metal-binding preferences. Predictions of small molecule structures using genome mining strategies have improved tremendously in recent years^18,19^; however, it is still not possible to predict the selectivity and affinity of metal coordination sites, and it is even difficult to predict whether a small molecule contains a metal-binding site^20,21^. This is because small molecule metal-binding sites are diverse and not conserved; as such, metal binding must be experimentally established. Various analytical methods including inductively coupled plasma mass spectrometry (ICP-MS)^22^, atomic absorption spectroscopy (AAS)^23^, x-ray fluorescence spectroscopy (XRF)^24^, UV-visible (UV-vis) absorption spectroscopy and nuclear magnetic resonance (NMR) spectroscopy in addition to multimodal approaches such as high-performance matrix-assisted laser desorption/ionization Fourier transform ion cyclotron resonance imaging mass spectrometry (MALDI FT-ICR IMS)^25^ can be used to analyze metal content and/or metal coordination. With each of these methods, it is still difficult to understand which molecules form metal complexes within a biological matrix that contains a pool of candidate metal ions, especially when the identities of metal-binding species are not known ahead of time. To address this shortcoming, we set out to develop a non-targeted liquid chromatography-tandem mass spectrometry (LC-MS/MS) based approach.

Screening for metal-binding compounds from complex biological matrices by non-targeted LC-MS/MS requires specific conditions to be met. Typical conditions used during sample extraction, preparation, and chromatography for non-targeted mass spectrometry involve low pH and high percentages of organic solvent, which both disfavor metal complexation. Even when a metal-bound species is observed, it commonly elutes at a different retention time than the apo (unbound) species and typically differs significantly in its MS/MS fragmentation behavior than the apo species. This inherent problem of conserving metal-ionophore complexes, in addition to differing retention times and differing gas phase fragmentation behavior (MS/MS) between apo- and metal-bound forms make it difficult to connect these forms to identify a given (metal specific) mass (*m/z*) offset. In order to overcome these limitations, we developed a two-step native electrospray ionization (ESI) MS/MS workflow, in which post-column metal infusion is coupled with pH adjustment via a double barrel syringe pump (**Figure 1a**) to directly detect candidate small molecules that bind to metals.

**Figure 1.**
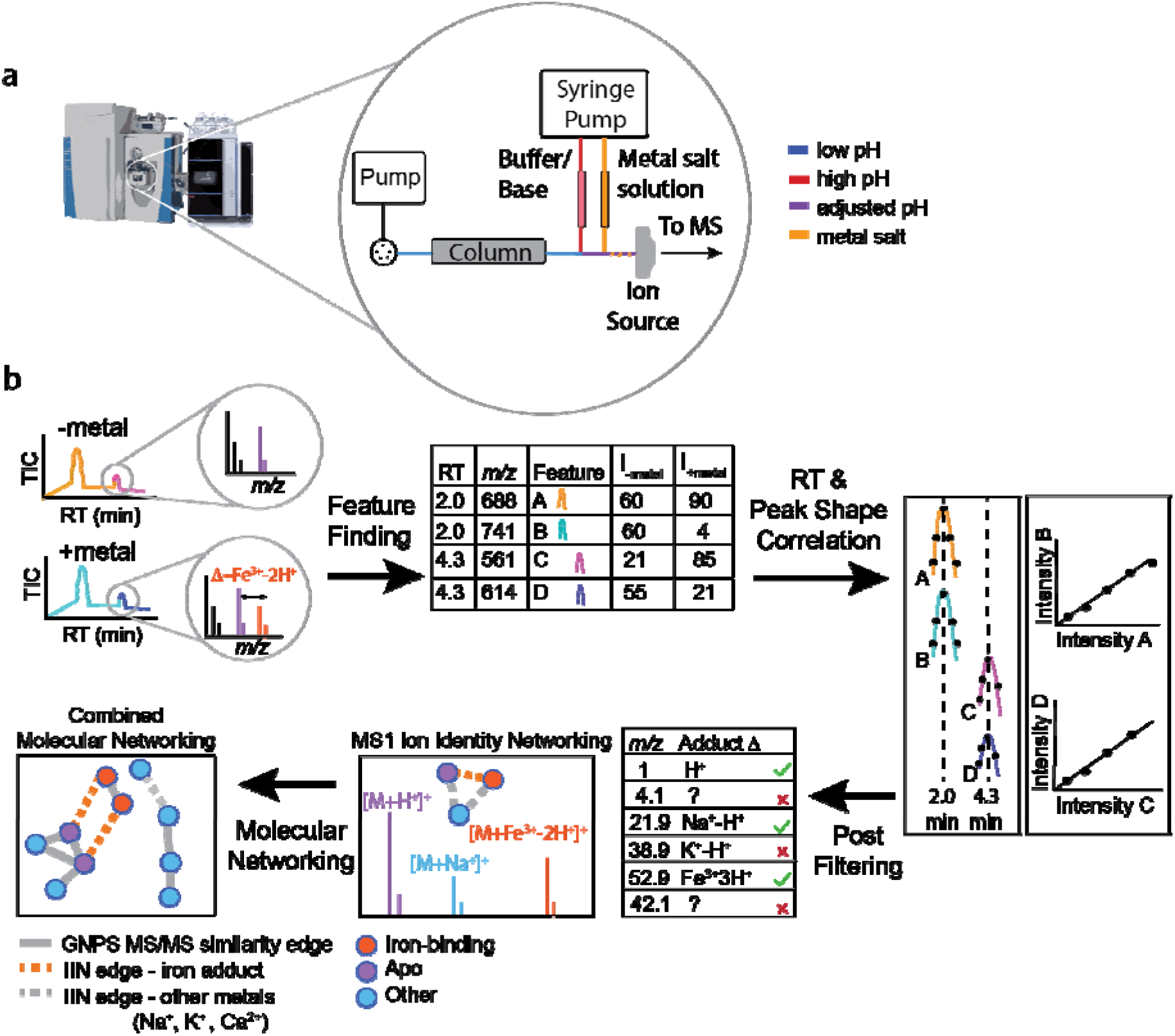
Overview of the native spray small molecule binding experiment. (a) The two-step native ESI-MS/MS workflow utilizes a post-column infusion of ammonium hydroxide solution followed by infusion of metal salt solution. (b) Data can be analyzed using a computational ion identity molecular networking (IIN) workflow in MZmine and GNPS.

After the implementation of post-column metal infusion that re-forms metal-ionophore complexes, one of the key challenges in developing this method was how to identify these metal-binding complexes within large datasets and/or complex mixtures that typically contain thousands of features. To overcome this limitation, a rule-based informatic workflow called ion identity molecular networking (IIN) was utilized (**Figure 1b**). Native ESI metal infusion coupled with IIN has enabled the systematic detection of candidate metal-binding molecules and metal-binding properties in different samples, such as bacterial and fungal culture extracts.

## Results

### Proof-of-Principle

Mass spectrometry-based methods for identifying metal-binding compounds using non-targeted metabolomics remain underexplored. We hypothesized that this is partially due to the conditions that disfavor metal coordination associated with sample preparation and separation; specifically, extractions often utilize high percentages of organic solvent, and reversed-phase LC separations are typically performed at low pH. Importantly, these conditions can alter both the configurations and ionization state of the molecules in solution, which can disrupt non-covalent interactions such as metal-binding.

To assess whether adjusting pH to physiological values, an approach often applied in intact protein mass spectrometry (native spray)^26–28^, would help with metal-ligand stability and detection, we incubated a series of ionophores with metals in buffered solutions at Ph 6.8 before direct infusion ESI-MS. Experiments were performed on a panel of siderophores, including ferrichrome (**1**), enterobactin (**2**), yersiniabactin (**3**), and desferrioxamine B (**4**). Each ionophore was added to a buffered solution at pH 6.8 in the presence or absence of iron(III) chloride salt (Fe^3+^). Direct infusion ESI-MS of these samples yielded predominantly apo (unbound) peaks in the absence of Fe^3+^ (**Figure 2a,c** and **SI Figure 1**) while peaks corresponding to both apo- (unbound) and iron-bound compound are observed in the presence of Fe^3+^ (**Figure 2b, d** and **SI Figure 1**). Peaks corresponding to apo and Fe^3+^-bound compound have an *m/z* delta of 52.9115, when comparing the iron-bound [M+Fe^3+^-2H^+^]^+^ and the apo [M+H^+^]^+^ species, where M is the neutral mass of the siderophore. pH-dependence of metal-binding is both compound and metal dependent^29,30^. Because both apo- and metal-bound species are observed at pH 6.8, which is a pre-requisite to observe the characteristic *m/z* delta of 52.9115, LC-MS/MS experiments were performed at this pH.

**Figure 2.**
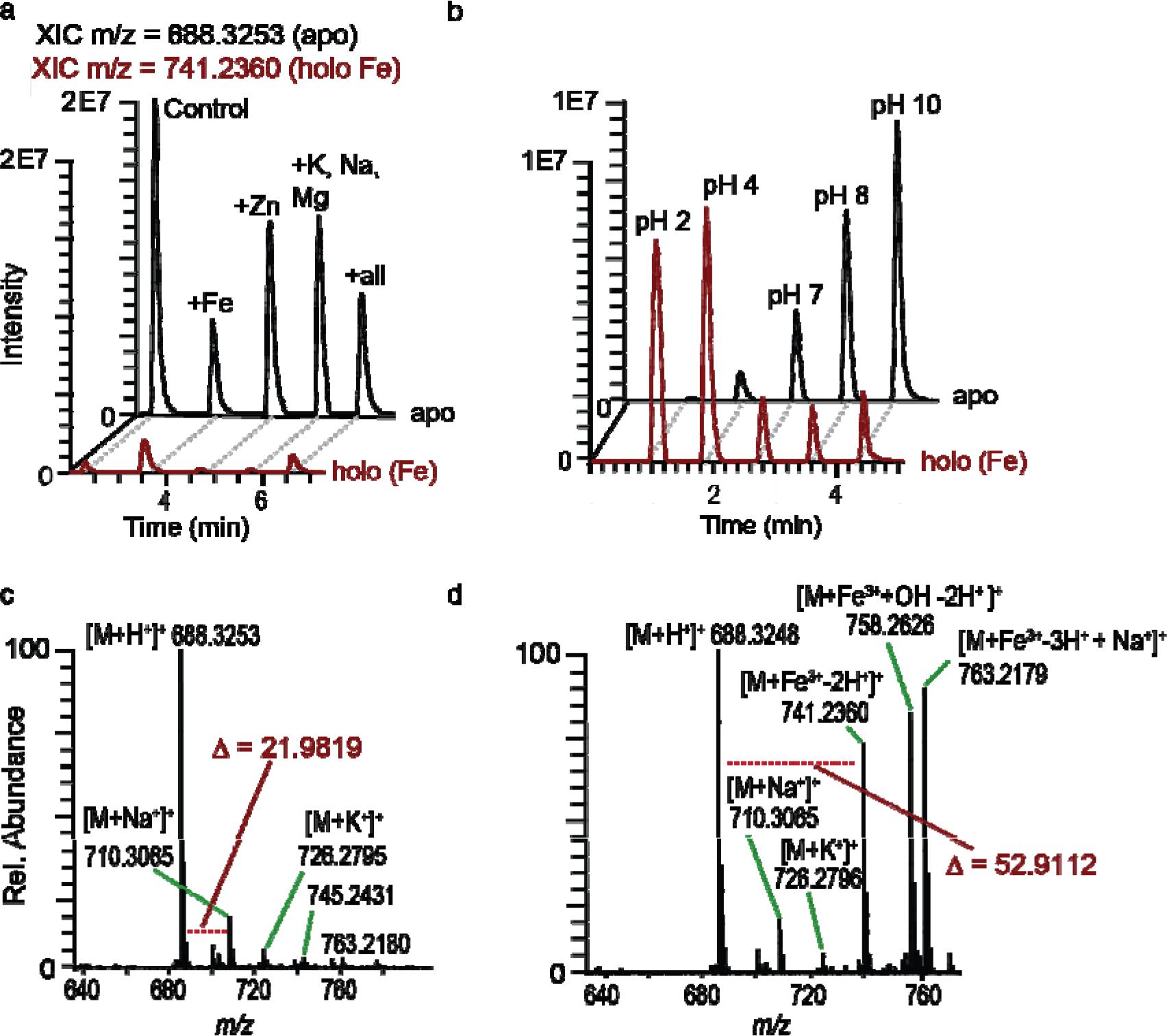
Direct infusion MS experiments on ferrichrome. In (a)-(b), each peak represents the direct injection of sample. The black trace represents the extracted ion chromatogram (XIC) of apo-ferrichrome ([M+H^+^]^+^, *m/z* = 688.32), while the red trace represents the extracted ion chromatogram (XIC) of Fe^3+^-bound ferrichrome ([M+Fe^3+^-2H^+^]^+^, *m/z* = 741.23). (a) Metal selectivity of binding was tested by adding a variety of metals (transition metals in addition to alkali and alkaline earth metals), and the iron-adduct is observed when Fe^3+^ is added in the absence and presence of other metals (“all” refers to the combination of Fe^3+^, Zn^2+^, K^+^, Na^+^, and Mg^2+^) but not when other metals are added in the absence of Fe^3+^. (b) pH-dependence of Fe^3+^-binding was assessed by performing a pH titration of ferrichrome in the presence of Fe^3+^. Direct infusion of ferrichrome in the presence of Fe^3+^ at pH 7 shows a nearly equal proportion of apo- to Fe^3+^-bound adduct. While (c) negligible Fe^3+^-bound peak is observed in the MS^1^ when apo-ferrichrome is injected at pH 7, (d) both apo- and Fe^3+^-bound peaks are observed when ferrichrome in the presence of Fe^3+^ is subjected to direct infusion MS. Apo and Fe^3+^-bound ferrichrome peaks are offset by specific *m/z* deltas.

Next, we found that metal binding is selective when a mixture of metal salts at equal ratios, including iron (III) chloride (Fe^3+^), zinc (II) acetate (Zn^2+^), and cobalt (II) chloride (Co^2+^), is added. While the Fe^3+^-bound species is the major form present when this metal mixture is added to ferrichrome (**1**), schizokinen (**5**), and desferrioxamine E (**6**) at pH 6.8, the Zn^2+^-bound species is the major metal-species present when the metal mixture is added to yersiniabactin (3) (**SI Figure 2**). It should be noted that a low (and pH dependent) intensity of Fe^3+^, Cu^+^, and Zn^2+^ ions was observed during analysis of the commercial yersiniabactin (**3**) in the absence of metals. Furthermore, direct infusion of Zn^2+^ to yersiniabactin was performed at various pH values – basic pH favors Zn^2+^-coordination^31^, which provided additional rationale for developing a modular LC-MS/MS workflow that could be run at pH 6.8. Recent work proposed that different products of the yersiniabactin operon may have multiple roles in biology. First, the yersiniabactin was shown to bind both iron and copper during urinary tract infections^32^; moreover, the yersiniabactin operon was shown to contribute to zinc acquisition by *Yersinia pestis* through an unknown mechanism^33^. Based on the above observations and others, we reassessed the yersiniabactin biology using genetic approaches in mouse models^31^. Taken as a whole, these evaluations suggest that native spray metal infusion could be a promising tool for facilitating the direct observation of metal complexes by ESI-MS.

Extracts from biological samples, however, are mixtures of hundreds to thousands of molecules; thus, it may not be optimal to analyze them using direct infusion. One strategy to de-complex such extracts is via the use of chromatographic separation. We therefore adapted the pH-adjusted direct infusion experiments to a LC-based workflow via a post-column infusion of ammonium hydroxide solution followed by metal salt solution in tandem (**Figure 1a**). As metal binding kinetics are fast (on the order of milliseconds^34–37^), we hypothesized that our set-up would allow for sufficient time to establish metal-ligand complexes after chromatography post pH adjustment and metal infusion, as it takes 0.6 seconds to reach the entrance of the electrospray inlet. Additionally, it is possible to delay the entrance to the instrument by lengthening the peek tubing post-metal infusion. Because metal complexes introduce a *m/z* shift, metal-binding is determined by observing a characteristic *m/z* delta between peaks corresponding to apo and metal-bound species with a similar chromatographic peak shape and retention time. However, as one LC-MS/MS run of a typical biological sample can contain hundreds to thousands of ion features, searching for the mass shifts associated with small molecule-metal interactions becomes challenging and tedious; based on this, we envisioned utilizing a computational workflow.

### Computational Workflow

Using this metal-infusion native ESI method to analyze a complex biological sample when one (or multiple) metals are infused post-LC presents a complex combinatorial problem of possible metal-small molecule binding interactions. A computational workflow was required to solve this; toward this end, we used ion identity molecular networking (IIN, **Figure 1b**)^38^ within the software tools MZmine 2^39,40^ linked with Global Natural Products Social Molecular Networking (GNPS)^41,42^. In IIN, LC-MS features, defined here as chromatographic peaks with a specific *m/z*, are grouped based on their retention time and chromatographic feature shape correlation and identified as specific ion types of the same analyte molecule akin to the way it is accomplished by CAMERA^43^ or RAMClust^44^. In this process, all metal-binding small molecules can be identified by linking the metal-bound ions to other adducts and in-source fragments of the apo structures. The connected ion features are then further linked within the molecular network created based on MS/MS similarity. As such, IIN is performed by (1) feature finding in MZmine 2^39,40^, (2) grouping of co-occurring features using Pearson correlation of their corresponding intensity profiles, (3) filtering results based on user-defined ion rules and *m/z* error tolerances, and (4) connection of similar MS/MS spectra through the cosine similarity metric. This enables the visualization of apo versus metal-bound compounds, even when the corresponding ion adducts have different gas phase fragmentation behavior in CID (**SI Figure 3**).

Toward this end, we envisioned comparing LC-MS/MS experiments with and without Fe^3+^-infusion and post-pH-adjustment. After performing feature finding that considers feature shapes in MZmine 2^39,40^, all grouped features are searched in pairs against a user-defined ion identity library, which is specified by lists of adducts, in-source fragments, and multimers. This approach tests all possible combinations of ion identities that point to the same neutral mass of a siderophore molecule (M). Consequently, as apo and bound-species have the same RT and feature shape in this workflow, IIN enables the discovery of novel metal-binding molecules by any combination of singly or multiply charged ion species and therefore lifts any restrictions to the most common ion types. This computational workflow facilitates the search for any characteristic *m/z* delta, defined by the user, to enable finding molecules that form complexes with iron (3+ ion, m/z delta=52.9115), in addition to other metals such as zinc (2+ ion, mass delta = 62.92077 Da), copper (2+ ion, 61.9212 Da), cobalt (2+ ion, 57.9248 Da), sodium (1+, 22.9892 Da), potassium (1+, 38.9632 Da), calcium (2+, 39.9615 Da), aluminum (3+, 24.9653 Da) etc.

### Assessing specificity and selectivity of post LC-infusion native spray MS in conjunction with IIN

In order to evaluate the computational workflow described above, we made three standard mixtures (corresponding to **Figure 3a, b**, and **c**, respectively) with molecules that are commercially available. When the siderophores yersiniabactin (**1**), vibriobactin (**2**), enterobactin (**3**), ferrichrome (**7**), and rhodotorulic acid (**6**) were subjected to post-LC pH adjustment and Fe^3+^-infusion, we observe the Fe^3+^-bound adducts of each siderophore; the Fe^3+^-bound siderophore is only observed post Fe^3+^-infusion (**Figure 3a**).

**Figure 3.**
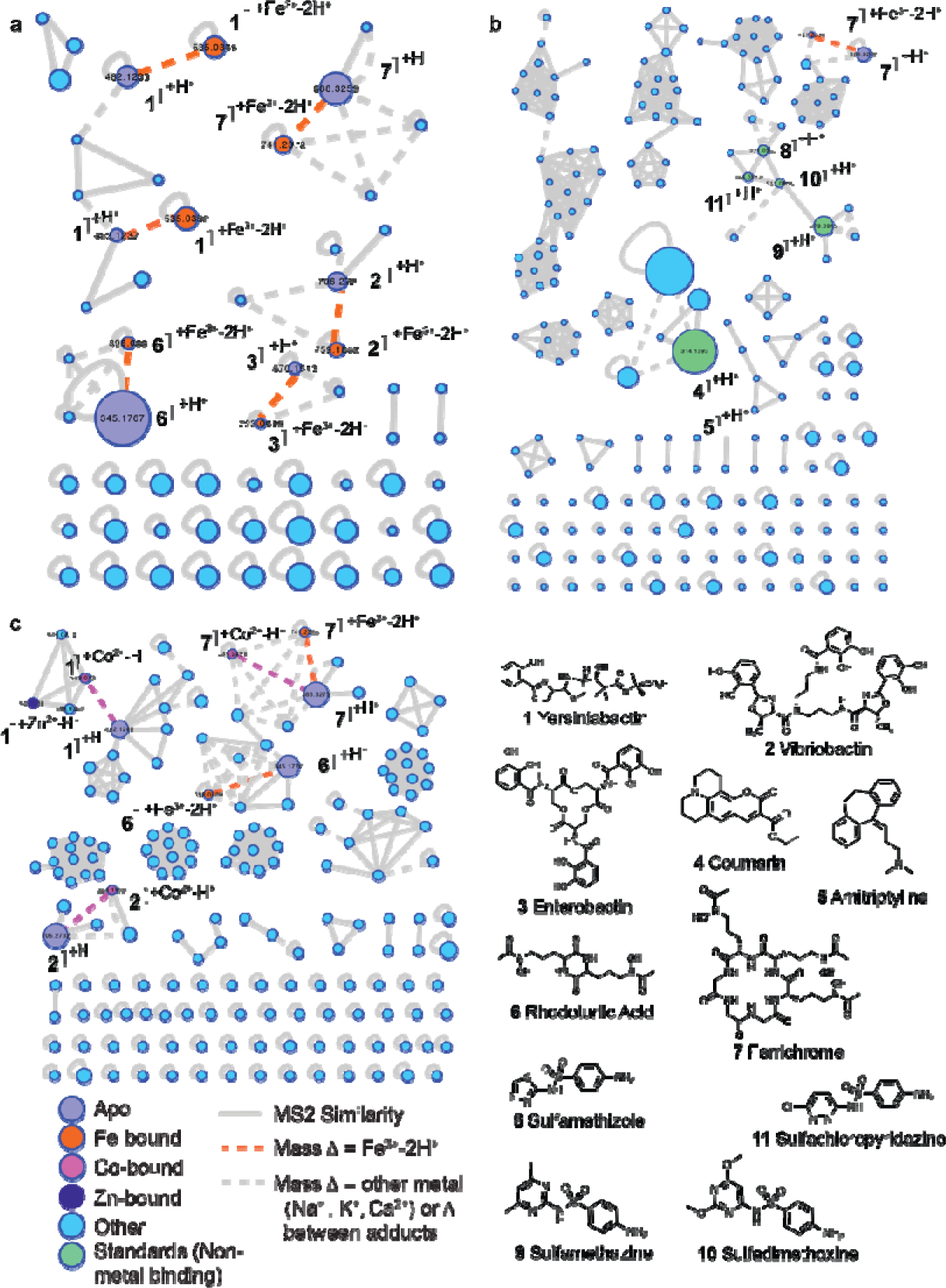
Post LC metal-infusion into mixtures of standards. (a) Post-LC infusion of Fe^3+^ and post-pH adjustment of a mixture containing ionophore standards **1,2,3,6** and **7**. Each of the six ionophore standards is complexed with iron (orange dashed line). (b) Post-LC infusion of Fe^3+^ and post-pH adjustment can be utilized to selectively observe the iron-bound adduct ferrichrome (**7**) in the presence of molecules **4,5,8,9,10**, and **11**, that have no reported iron-binding behavior. (c) Ionophores **1,2,6** and **7** form adducts when subjected to native spray with a mixture of metals after post-LC infusion. The mixture of metals includes Co^2+^, Fe^3+^, and Zn^2+^. Nodes corresponding to an [M+H^+^]^+^ ion of apo siderophores are colored in purple and nodes corresponding to iron-bound siderophores are colored in orange. MS/MS similarity edges are solid grey lines, IIN edges corresponding to an iron *m/z* delta are dashed orange lines, and IIN edges corresponding to other metals (such as Na^+^, Ca^2+^, and K^+^) or *m/z* deltas between different combinations of metals. The size of the node corresponds to relative size of the feature.

To assess whether metal-binding exhibits specificity or whether non-specific adducts form after post-LC Fe^3+^-infusion, we added competing molecules – molecules without known Fe^3+^-binding properties – to ferrichrome. We then analyzed this mixture using LC-MS/MS with and without pH adjustment and Fe^3+^-infusion. In this experiment, ferrichrome is the only compound connected with an Fe^3+^-binding edge and no other Fe^3+^-binding to other molecules are observed (**Figure 3b**). To further assess the selectivity of the metal binding, four siderophores were subjected to a cocktail of metals (Zn^2+^, Fe^3+^, and Co^2+^) after chromatography and pH adjustment to 6.8 (**Figure 3c**). Using IIN, we searched for Zn^2+^, Fe^3+^, Co^2+^, Na^+^, K^+^, and Ca^2+^ adducts. This revealed that all the non-ionophore molecules added to the sample did not have any metal adducts, while all the ionophores did. It further revealed specificity by the different ionophores. For example, yersiniabactin (**1**) preferentially binds divalent metals at this pH, which is consistent with our investigation of yersiniabactin biology^31^ in addition to prior studies^32,45^. Rhodotorulic acid (**6**) exhibits a strong selectivity for Fe^3+^, ferrichrome (**7**) is less specific, exhibiting both Fe^3+^- and Co^2+^-binding, while vibriobactin (**2**) shows only Co^2+^-binding. This data suggests that each ionophore has preferences for binding specific metals, otherwise adducts of each of the three metals would have been observed in equal proportions.

### Observing Siderophores in Bacterial Culture Extracts

In order to further assess selectivity in the presence of non-binding compounds found in biology, we tested this workflow on supernatant extracts from *Glutamicibacter arilaitensis. JB182*^46^. This microbe was previously isolated from cheese and grown here in liquid cheese media. Cheese is an iron-depleted environment, as it contains lactoferrin, an iron-binding protein that sequesters free iron^48^. Siderophore production was predicted based on genomic data mining using antiSMASH 4.0^47^. From the genome, antiSMASH^47^ predicted the presence of biosynthetic gene clusters for ferrioxamine B and fuscachelin with 50% and 44% similarity, respectively. Using the native metal metabolomics method developed here, both apo-desferrioxamine E (DFE) and the Fe^3+^-bound ferrioxamine E were observed from culture extracts and were connected by an Fe^3+^-binding IIN edge. DFE was annotated as a spectral match provided via molecular networking in GNPS^41,42^. Strikingly, DFE was the only Fe^3+^-binding connection observed using IIN (**Figure 4a**); this important observation illustrates the specificity of Fe^3+^-binding during the post-LC infusion as Fe^3+^ binds only one specific molecule and none of the other molecules detected in this complex biological sample (**Figure 4b-e**).

**Figure 4.**
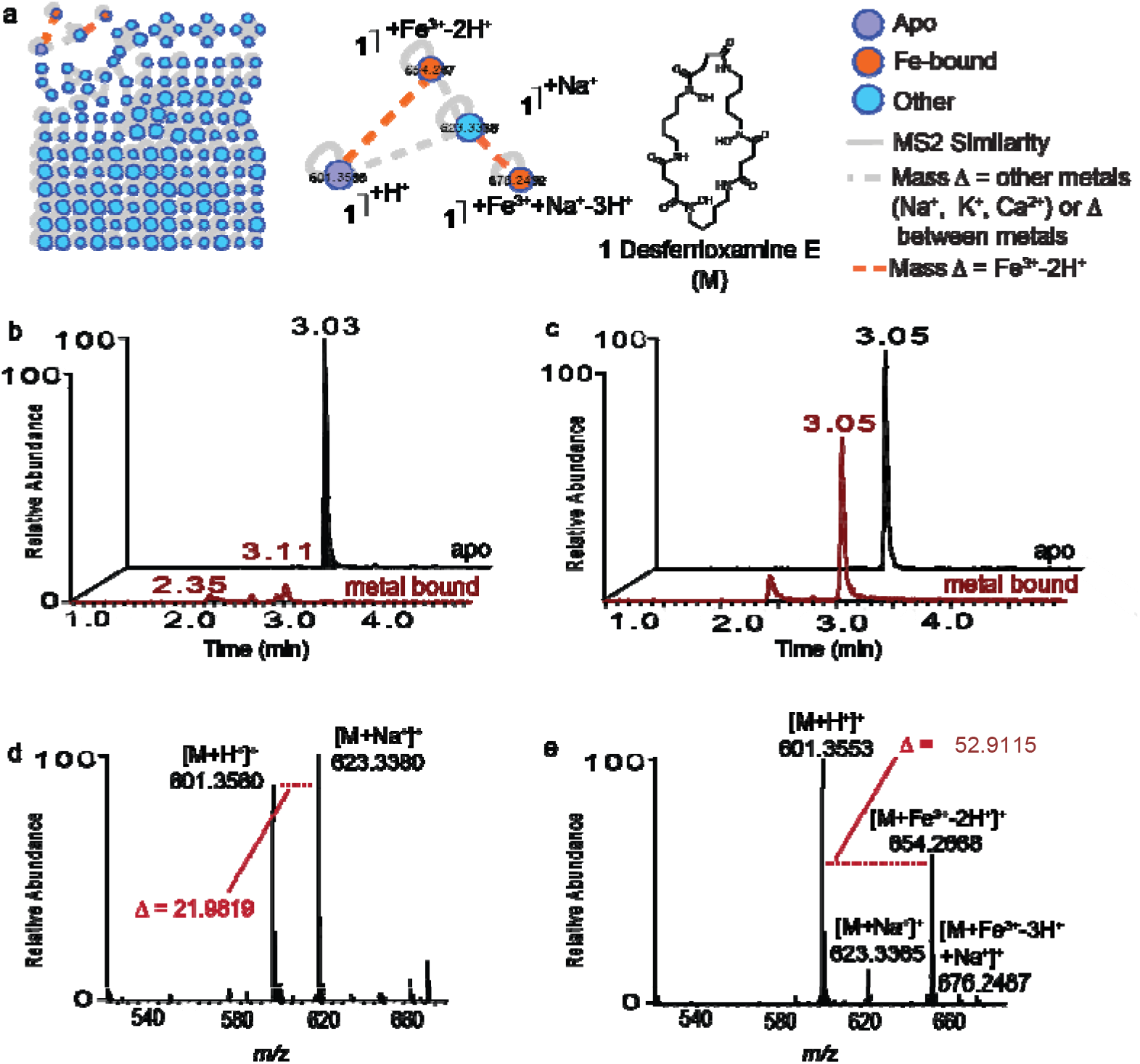
Native spray metal metabolomics is used to identify siderophores in bacterial culture extracts. (a) Desferrioxamine E (DFE) is identified as the only siderophore present in *Glutamicibacter arilaitensis. JB182* extracts. The only Fe^3+^-binding IIN edges observed in the full network correspond to DFE and the sodium adduct of DFE. (b) Extracted Ion Chromatogram of apo- DFE (*m/z* 601. 35, black) and Fe^3+^-bound DFE (*m/z* 654.26, red) when analysis is performed using standard LC-MS/MS conditions. (c) Extracted Ion Chromatogram of apo-DFE (*m/z* 601. 35, black) and Fe^3+^-bound DFE (*m/z* 654.26, red) when analysis is performed using Fe^3+^-infusion LC-MS/MS conditions. (d) MS^1^ of the DFE peak (at retention time 3.03 min) in (b). Only the apo- and Na^+^-adduct are observed, with a *m/z* delta (Δ) of 21.9819, but no Fe^3+^-bound peak is observed. (e) MS^1^ of the DFE peak (at retention time 3.05 min) in (c). The apo- and the Fe^3+^-bound peak are observed, with a m/z delta (Δ) of 53 9115

Based on the selectivity experiments with commercially available yersiniabactin, we were also interested in further investigating an organism that produces yersiniabactin in addition to other siderophores in culture. The probiotic *Escherichia coli* strain Nissle 1917 (*E. coli* Nissle, serotype O6:K5:H1) is known to produce yersiniabactin^49^, and we hypothesized that we could use the native spray strategy to observe ionophore metal-binding from an *E. coli* Nissle supernatant extract grown in M9 glucose minimal medium. We therefore ran the extract with and without metal infusion and found a number of iron binding molecules using IIN (**Figure 5a** and **SI Figure 4**). From this culture we detected yersiniabactin, aerobactin, and HPTzTn-COOH^50^, a truncated derivative of yersiniabactin, and each corresponding Fe^3+^-complex (**Figure 5b-d**). IIN suggests a number of additional yersiniabactin (**Figure 5b and c**) and aerobactin (**Figure 5d**) derivatives that also bind iron. Using SIRIUS 4.0^1^ revealed molecular formulas, suggesting at least fifteen additional siderophores are present in this culture extract. These siderophores that co-network are structurally related to the three known siderophores and their predicted molecular formulas are tabulated in Table 1a. This is consistent with the observation that many natural product gene clusters make many related molecules^51^

**Table 1.**
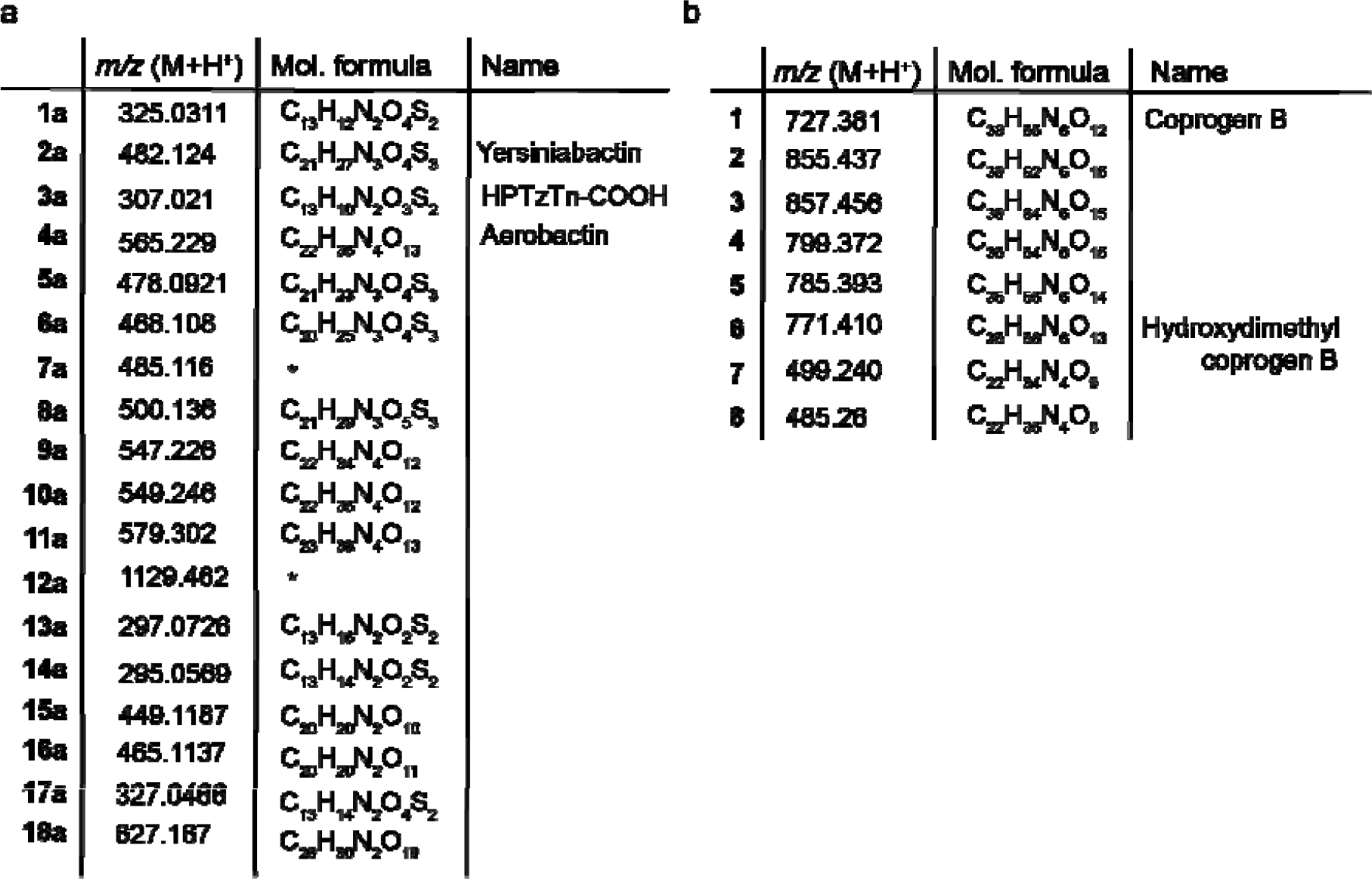
Iron-binding derivatives identified in *E. coli* Nissle (a) and *Eutypa lata* (b) cultures have been tabulated above. The molecular (mol.) formula corresponds to the formula with the top score assigned in SIRIUS 4.0^1^, and those compounds with less than 1 matched peaks in SIRIUS 4.0 have been labeled with an asterisk (*).

**Figure 5.**
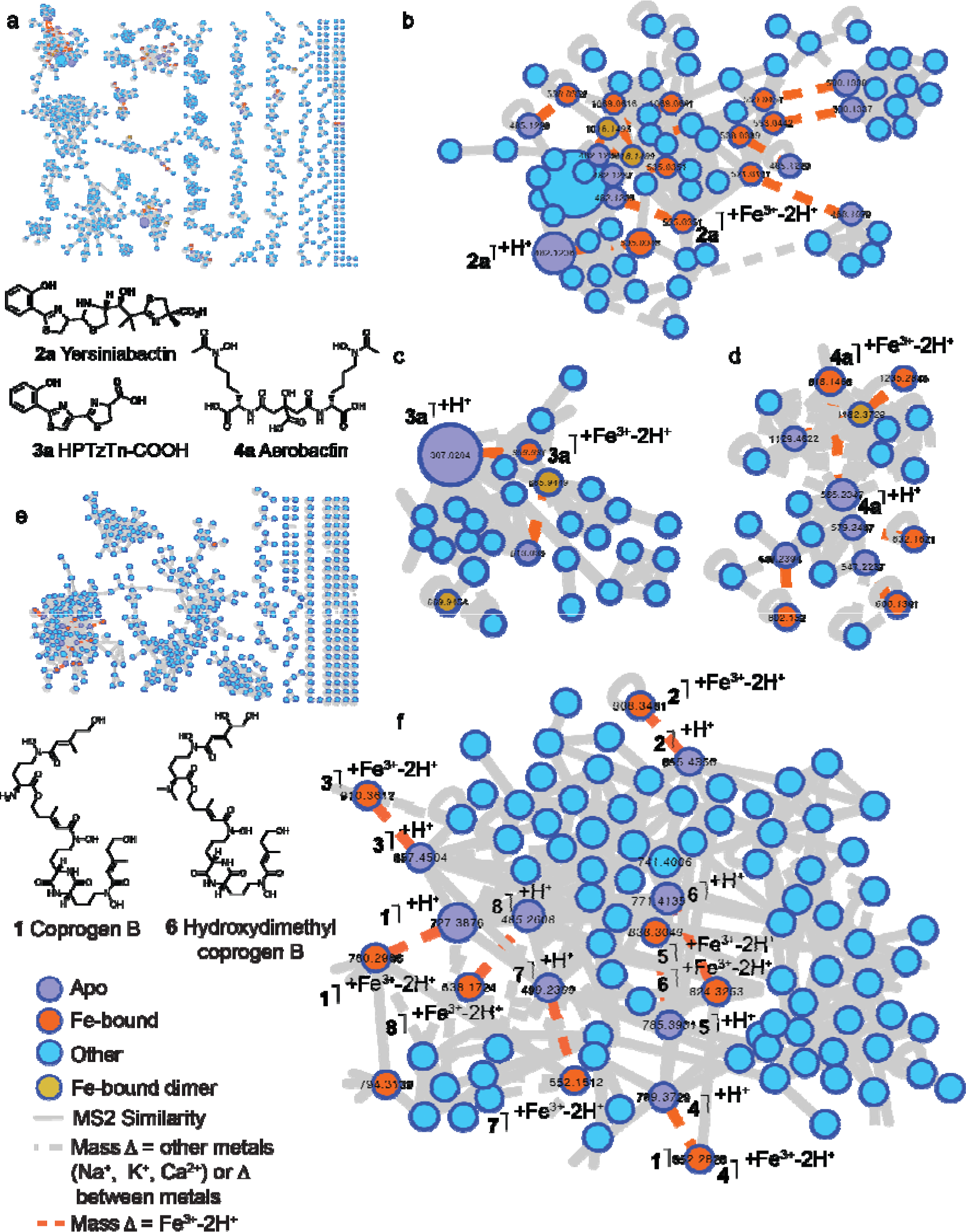
Native spray metal metabolomics is used to identify known and novel siderophores in bacterial and fungal culture extracts. (a) A number of siderophores are observed in *E. coli* Nissle culture extracts when a post-LC infusion of Fe^3+^ and post-pH adjustment is performed. Singletons have been removed from this network. A number of (b) yersiniabactin (**2a**) derivatives, including (c) the truncated biosynthetic intermediate of yersiniabactin, HPTzTn (**3a**), in addition to (h) aerobactin (**4a**) derivatives are identified using native metal metabolomics. These derivatives are tabulated in Table 1a. (e) A number of siderophores are also observed in *Eutypa lata* culture extracts when a post-LC infusion of Fe^3+^ and base are performed, (f) corresponding to a number of coprogen B (**1**) and hydroxydimethyl coprogen B (**6**) derivatives (**2-5**, **7**-**8**), which are tabulated in Table 1b.

### Observing Meta I Binding Molecules from a Eukaryote

After illustrating the ability of native spray metal meta bolomics to specifically elucidate metal-binding molecules present in *E. coli* Nissle extracts, we next tested this method on increasingly complex samples. Toward this end, we applied our method to fungal extracts; specifically, we chose to analyze extracts from the wine fungus *Eutypa lata*. The *Eutypa lata* genome^52^ is approximately ten times the size of the *E. coli* Nissle genome^53^. Moreover, the published *Eutypa lata* strain UCREL1 contains a predicted domain structure that matches the coprogen synthetase SSM2 from *Magnaporthe oryzae*^54–56^. Applying the post-LC pH neutralization and Fe^3+^-infusion mass spectrometry workflow enabled detection of apo-coprogen B and apo-hydroxydimethyl coprogen B in addition to the iron-bound forms of both siderophores. Apo and Fe^3+^-bound forms were each connected through an Fe^3+^-binding IIN edge (**Figure 5e**). We additionally identified a number of structurally related derivatives in the same molecular family that bind iron (**Figure 5f**). As in the *E. coli* Nissle example, we used SIRIUS 4.0^1^ to elucidate molecular formulas and have putatively annotated six structurally related novel siderophores, again underscoring the potential of this method for discovery. These siderophores are tabulated in Table 1b.

## Conclusion

The native ESI and post-LC metal-infusion workflow described here facilitates the characterization of known and novel ionophores. As this native metal metabolomic approach is simple to implement on any liquid chromatography-based mass spectrometry system, we anticipate this method can become a useful and widespread strategy for elucidating metal-binding small molecules in biology. Currently, analysis of metal-binding is difficult to perform on complex samples, as the majority of techniques used to assess metal binding (including NMR, UV-vis, EPR) are usually performed on purified compounds to obtain meaningful results. ICP-MS and AAS can be used to assess bulk metal content of complex samples; however, they are not able to provide molecular detail about which compounds present in a complex sample actually bind metal; split flow LC systems^21^ provide innovative strategies for simultaneously analyzing metal content and structures but require expensive and specialized equipment and customized set-ups. Based on ease of implementation, we foresee routine implementation of this native metal metabolomic approach into non-targeted LC-MS/MS metabolomics workflows. Additionally, this workflow can be adapted to give detailed insights into metal-binding selectivity, and we hypothesize this method could be used to estimate relative metal-binding affinities of small molecules. Based on our results, we suspect that molecules traditionally described as siderophores may also be responsible for binding and uptake of other metals; yersiniabactin is one example of this^31^, as we provide direct evidence that it binds to zinc, in addition to copper and iron. The method outlined here is uniquely set up to identify novel metal-binding molecules in complex samples and we anticipate that methods that enable their detection will significantly improve our understanding of the role of metals across diverse fields ranging from human health to agriculture to the environment.

## Online Methods

### Sample Preparation

Commercial ionophore standards were purchased from EMC Microcollections, https://www.microcollections.de/product-services/siderophores.html. Standards were diluted in either water or methanol to a concentration of 1 mM and diluted further from the stock solution.

### Direct Inject-MS data acquisition

For MS analysis, 5 μL were directly infused into a Q-Exactive orbitrap mass spectrometer via flow injections through a Vanquish UHPLC system (Thermo Fisher Scientific, Bremen, Germany). A flow rate between 0.2 mL/min and 0.4 mL/min was used. MS^1^ data acquisition was performed in positive mode. Electrospray ionization (ESI) parameters were set to 53 L/min sheath gas flow, 14 L/min auxiliary gas flow, 0 L/min sweep gas flow and 400°C auxiliary gas temperature. The spray voltage was set to 3.5 kV and the inlet capillary to 320°C. 50 V S-lens level was applied. MS scan range was set to 150-1500 m/z with a resolution at m/z 200 (R_m/z 200_) of 35,000 with one micro-scan. The maximum ion injection time was set to 100 ms with an automated gain control (AGC) target of 1.0E6.

### LC-MS/MS data acquisition

For LC-MS/MS analysis, 2-5 μL were injected into a Vanquish UHPLC system coupled to a Q-Exactive orbitrap mass spectrometer (Thermo Fisher Scientific, Bremen, Germany). For the chromatographic separation, a C18 porous core column (Kinetex C18, 50 × 2 mm, 1.8 um particle size, 100 A pore size, Phenomenex, Torrance, USA) was used. For gradient elution, a high-pressure binary gradient system was used. The mobile phase consisted of solvent A H_2_O + 0.1 % formic acid (FA) and solvent B acetonitrile (ACN) + 0.1 % FA, unless otherwise specified. The flow rate was set to 0.5 mL/min, unless otherwise specified. One of the following two methods (*method 1* or *method 2*) was used for analysis. After injection, the samples were eluted with one of the following linear gradients: 0-0.5 min, 5% B, 0.5-5 min 5-99% B, followed by a 2 min washout phase at 99% B and a 3 min re-equilibration phase at 5% B (*method 1*) or 0-0.5 min, 5 % B, 0.5-9 min 5-100% B, followed by a 2 min washout phase at 99% B and a 3 min re-equilibration phase at 5% B. (*method 2*). Data dependent acquisition (DDA) of MS/MS spectra was performed in positive mode. Electrospray ionization (ESI) parameters were set to 53 L/min sheath gas flow, 14 L/min auxiliary gas flow, 0 L/min sweep gas flow and 400°C auxiliary gas temperature; the spray voltage was set to 3.5 kV and the inlet capillary to 320°C and 50 V S-lens level was applied (*method 1*) or 60 L/min sheath gas flow, 20 L/min auxiliary gas flow, 3 L/min sweep gas flow and 300°C auxiliary gas temperature; the spray voltage was set to 3.5 kV and the inlet capillary to 380°C. 60 V S-lens level was applied (*method 2*). MS scan range was set to 150-1500 m/z with a resolution at m/z 200 (R_m/z 200_) of 35,000 with one micro-scan. The maximum ion injection time was set to 100 ms with an automated gain control (AGC) target of 1.0E6. Up to 5 MS/MS spectra per MS1 survey scan were recorded DDA mode with R_m/z 200_ of 17,500 with one micro-scan. The maximum ion injection time for MS/MS scans was set to 100 ms with an AGC target of 1E6 ions (*method 1*) or 5E5 ions (*method 2*). The MS/MS precursor isolation window was set to m/z 1 (*method 1*) or 2 (*method 2*). Normalized collision energy was set to a stepwise increase from 20 to 30 to 40% with z = 1 as default charge state. MS/MS scans were triggered at the apex of chromatographic peaks within 2 to 15 s from their first occurrence. Dynamic precursor exclusion was set to 5 s (*method 1*) or 30 s (*method 2*). Ions with unassigned charge states were excluded from MS/MS acquisition as well as isotope peaks. *Method 2:*

### Data analysis

Feature finding and ion identity networking was performed using an in-house modified version of MZmine2.37^38–40^, corr.17.7 available at https://github.com/robinschmid/mzmine2/releases. Feature tables, MS/MS spectra files (mgf), and ion identity networking results were exported, uploaded to MassIVE, and submitted to GNPS for feature-based molecular networking analysis. Details of analysis are provided in **Supplementary Methods**.

### Data and software availability

All mass spectrometry .raw and centroided .mzXML or .mzML files are publicly available in the mass spectrometry interactive virtual environment (MassIVE) under massive.ucsd.edu with project identifier <u>MSV000084237</u> (Standards), <u>MSV000084289</u> (Cheese siderophores), <u>MSV000082999</u> (Fungal siderophores), <u>MSV000083387</u> (*E.coli* Nissle siderophores). Ion Identity Molecular Networks (IIN) can be accessed through gnps.ucsd.edu under direct links: https://gnps.ucsd.edu/ProteoSAFe/status.jsp?task=79d0f380b4814ff9a720836c5570036f, https://gnps.ucsd.edu/ProteoSAFe/status.jsp?task=ad4b2665dfb744d09a9d2445f1213720, http://gnps.ucsd.edu/ProteoSAFe/status.jsp?task=0d8cacfd74744357ada2f7a081252d64, https://gnps.ucsd.edu/ProteoSAFe/status.jsp?task=5459d22126e843a3a1449f8362cd267f, http://gnps.ucsd.edu/ProteoSAFe/status.jsp?task=1c3e79f0ab984386bd468e2d163281e0, http://gnps.ucsd.edu/ProteoSAFe/status.jsp?task=256ba734f4334c1c90f65ffbd9141d0e.

The modified version of MZmine 2 (2.37, corr.17.7)^38–40^ can be found here https://github.com/robinschmid/mzmine2/releases. GNPS^41,42^ can be accessed here https://gnps.ucsd.edu/ProteoSAFe/static/gnps-splash.jsp Cytoscape^57^ version 3.7.1 can be accessed here https://cytoscape.org/.

## Acknowledgements

We want to thank the following sources for support P41-GM103484, GMS10RR029121 and the Betty and Gordon Moore Foundation. DP was supported through the Deutsche Forschungsgemeinschaft with grant PE 2600/1. Work in MR lab is supported by Public Health Service Grants AI126277, AI114625, AI145325, by the Chiba University-UCSD Center for Mucosal Immunology, Allergy, and Vaccines, and by the UCSD Department of Pediatrics. M.R. also holds an Investigator in the Pathogenesis of Infectious Disease Award from the Burroughs Wellcome Fund.

## References

1 Dührkop, K. et al. SIRIUS 4: a rapid tool for turning tandem mass spectra into metabolite structure information. Nat Methods 16, 299–302 (2019).

2 Holden, V. I. & Bachman, M. A. Diverging roles of bacterial siderophores during infection. Metallomics 7, 986–995 (2015).

3 Sandy, M. & Butler, A. Microbial Iron Acquisition: Marine and Terrestrial Siderophores. Chem Rev. 109, 4580–4595 (2009).

4 Raymond, K. N., Allred, B. E. & Sia, A. K. Coordination Chemistry of Microbial Iron Transport. Acc. Chem. Res. 48, 2496–2505 (2015).

5 Vraspir, J. M. & Butler, A. Chemistry of Marine Ligands and Siderophores. Annu. Rev. Mar. Sci 1, 43–63 (2009).

6 Kenney, G. E. & Rosenzweig, A. C. Chalkophores. Annu. Rev. Biochem. 87, 645–676 (2018).

7 Wang, L. et al. Diisonitrile Natural Product SF2768 Functions As a Chalkophore That Mediates Copper Acquisition in Streptomyces thioluteus. ACS Chem Biol. 12, 3067–3075 (2017).

8 Łobodaa, D. & Rowińska-Żyrek, M. Zinc binding sites in Pra1, a zincophore from Candida albicans. Dalton Trans. 46, 13695–13703 (2017).

9 Johnstone, T. C. & Nolan, E. M. Beyond Iron: Non-Classical Biological Functions of Bacterial Siderophores. Dalton Trans. 44, 6320–6339 (2015).

10 Wilson, D., Citiulo, F. & Hube, B. Zinc Exploitation by Pathogenic Fungi. PLOS Pathog 8, e1003034 (2012).

11 Capdevila, D. A., Wang, J. & Giedroc, D. P. Bacterial Strategies to Maintain Zinc Metallostasis at the Host-Pathogen Interface. J Biol Chem 291, 20858–20868 (2016).

12 Hover, B. M. et al. Culture-independent discovery of the malacidins as calcium-dependent antibiotics with activity against multidrug-resistant Gram-positive pathogens. Nat Microbiol 3, 415–422 (2018).

13 Wood, T. M. & Martin, N. I. The calcium-dependent lipopeptide antibiotics: structure, mechanism, & medicinal chemistry. Med. Chem. Commun. 10, 634–646 (2019).

14 D., J., A., R., M., O. & E., H. R. Structural transitions as determinants of the action of the calcium-dependent antibiotic daptomycin. Chem Biol 11, 949–957 (2004).

15 von Eckardstein, L. et al. Total Synthesis and Biological Assessment of Novel Albicidins Discovered by Mass Spectrometric Networking. Chem.: Eur. J. 23, 15316–15321 (2017).

16 Petras, D. et al. High-Resolution Liquid Chromatography Tandem Mass Spectrometry Enables Large Scale Molecular Characterization of Dissolved Organic Matter. Front. Mar. Sci., doi:10.3389/fmars.2017.00405 (2017).

17 McDonald, D. e. a. American Gut: an Open Platform for Citizen Science Microbiome Research. mSystems 3, e00031–00018, doi:10.1128/mSystems.00031-18 (2018).

18 Medema, M. H. & Fischbach, M. A. Computational approaches to natural product discovery. Nat Chem Biol. 11, 639–648 (2015).

19 Bachmann, B. O., Van Lanene, S. G. & Baltz, R. H. Microbial genome mining for accelerated natural products discovery: is a renaissance in the making? J Ind Microbiol Biotechnol 41, 175–184 (2014).

20 Kasampalidis, I. N., Pitas, I. & Lyroudia, K. Conservation of metal-coordinating residues. Proteins: Struct. Funct. Bioinf. 68, 123–130 (2007).

21 Cvetkovic, A. et al. Microbial metalloproteomes are largely uncharacterized. Nature 466, 779–782 (2010).

22 Ackerman, C. M., Lee, S. & Chang, C. J. Analytical Methods for Imaging Metals in Biology: From Transition Metal Metabolism to Transition Metal Signaling. Anal Chem 89, 22–41 (2017).

23 Piper, K. G. & Higgins, G. Estimation of Trace Metals in Biological Material by Atomic Absorption Spectrophotometry. Proc. Ass. Clin. Biochem. 4, 190–197 (1967).

24 Aschner, M. et al. Imaging metals in Caenorhabditis elegans. Metallomics 9, 357–364 (2017).

25 Perry, W. J. et al. Staphylococcus aureus exhibits heterogeneous siderophore production within the vertebrate host. Proc. Nat. Acad. Sci. USA, doi:10.1073/pnas.1913991116 (2019).

26 Leney, A. C. & Heck, A. J. R. Native Mass Spectrometry: What is in the Name? J Am Soc Mass Spectrom 28, 5–13 (2017).

27 Zhang, H., Cui, W. & Gross, M. L. Native electrospray ionization and electron-capture dissociation for comparison of protein structure in solution and the gas phase. Int J Mass Spectrom 354, 288–291 (2013).

28 Beveridge, R. et al. Mass spectrometry locates local and allosteric conformational changes that occur on cofactor binding. Nat Commun 7, 12163 (2016).

29 Miethke, M. & Marahiel, M. A. Siderophore-Based Iron Acquisition and Pathogen Control. Microbiol Mol Biol Rev. 71, 413–415 (2007).

30 Hider, R. C. & Kong, X. Chemistry and biology of siderophores. Nat Prod Rep 27, 637–657 (2009).

31 Zhi, H. et al. Probiotic Escherichia coli Nissle 1917 Uses Zinc transporters and the Siderophore Yersiniabactin to Acquire Zinc in the inflamed gut. draft in preparation (2019).

32 Chaturvedi, K. S., Hung, C. S., Crowley, J. R., Stapleton, A. E. & Henderson, J. P. The siderophore yersiniabactin binds copper to protect pathogens during infection. Nat Chem Biol 8, 731–736 (2012).

33 Bobrov, A. G. et al. The Yersinia pestis siderophore, yersiniabactin, and the ZnuABC system both contribute to zinc acquisition and the development of lethal septicaemic plague in mice. Mol Microbiol 93, 759–775 (2014).

34 Billo, E. J., Brito, K. K. & Wilkins, R. G. Kinetics of Formation and Dissociation of Metallocarboxypeptidases. Bioinorganic Chem 8, 461–475 (1978).

35 Magzoub, M., Padmawar, P., Dix, J. A. & Verkman, A. S. Millisecond association kinetics of K+ with triazacryptand-based K+ indicators measured by fluorescence correlation spectroscopy. J Phys Chem B. 110, 21216–21221 (2006).

36 Chock, P. B. Relaxation Study of Complex Formation between Monovalent Cations and Cyclic Polyethers. Proc. Nat. Acad. Sci. USA 69, 1939–1942 (1972).

37 Pasternack, R. F., Gipp, L. & Sigel, H. Thermodynamics and Kinetics of Complex Formation between Cobalt(II), Nickel(II), and Copper (II) with Glycyl-L-leucine and L-Leucylglycine. J Am Chem Soc 94, 8031–8038 (1972).

38 Schmid, R. et al. Ion identity based molecular networking. In preparation (2019).

39 Pluskal, T., Castillo, S., Villar-Briones, A. & Orešič, M. MZmine 2: Modular framework for processing, visualizing, and analyzing mass spectrometry-based molecular profile data. BMC Bioinformaticsvolume 11, 395 (2010).

40 Katajamaa, M., Miettinen, J. & Oresic, M. MZmine: toolbox for processing and visualization of mass spectrometry based molecular profile data. Bioinformatics 22, 634–636 (2).

41 Wang, M. Sharing and community curation of mass spectrometry data with Global Natural Products Social Molecular Networking. Nat Biotechnol. 34, 828–837 (2016).

42 Aron, A. T. et al. Reproducible Molecular Networking Of Untargeted Mass Spectrometry Data Using GNPS. ChemRxiv Preprint, doi:10.26434/chemrxiv.9333212.v1 (2019 ChemRxiv. Preprint).

43 Kuhl, C., Tautenhahn, R., Böttcherr, C., Larson, T. R. & Neumann, S. CAMERA: An integrated strategy for compound spectra extraction and annotation of LC/MS data sets. Anal Chem 84, 283–289 (2012).

44 Broeckling, C. D., Afsar, F. A., Neumann, S., Ben-Hur, A. & Prenni, J. E. RAMClust: A Novel Feature Clustering Method Enables SpectralMatching-Based Annotation for Metabolomics Data. Anal Chem 86, 6812–6817 (2014).

45 Robinson, A. E., Lowe, J. E., Koh, E. & Henderson, J. P. Uropathogenic enterobacteria use the yersiniabactin metallophore system to acquire nickel. J Biol Chem 293, 14953–14961 (2018).

46 Bonham, K. S., Wolfe, B. E. & Dutton, R. J. Extensive horizontal gene transfer in cheese-associated bacteria. eLife 6, e22144, doi:10.7554/eLife.22144 (2017).

47 Blin, K. et al. antiSMASH 4.0—improvements in chemistry prediction and gene cluster boundary identification. Nucleic Acids Res 45, W36–W41 (2017).

48 Monnet, C., Back, A. & Irlinger, F. Growth of aerobic ripening bacteria at the cheese surface is limited by the availability of iron. Appl Environ Microbiol. 78, 3185–3192 (2012).

49 Valdebenito, M., Crumbliss, A. L., Winkelmann, G. & K., H. Environmental factors influence the production of enterobactin, salmochelin, aerobactin, and yersiniabactin in Escherichia coli strain Nissle 1917. Int J Med Microbiol. 296, 513–520 (2006).

50 Xu, G., Guo, H. & Lv, H. Metabolomics Assay Identified a Novel Virulence-Associated Siderophore Encoded by the High-Pathogenicity Island in Uropathogenic Escherichia coli. J. Proteome Res. 18, 2331–2336 (2019).

51 Fischbach, M. A. & Clardy, J. One pathway, many products. Nat Chem Biol 3, 353–355 (2007).

52 Blanco-Ulate, B., Rolshausen, P. E. & Cantu, D. Draft Genome Sequence of the Grapevine Dieback Fungus Eutypa lata UCR-EL1. genome A 1, e00228–00213 (2013).

53 Reistera, M. et al. Complete genome sequence of the Gram-negative probiotic Escherichia coli strain Nissle 1917. J of Biotech 187, 106–107 (2014).

54 Hof, C., Eisfeld, K., Antelo, L., Foster, A. J. & Anke, H. Siderophore synthesis in Magnaporthe grisea is essential for vegetative growth, conidiation and resistance to oxidative stress. Fungal Genet Biol 46, 321–332 (2009).

55 Antelo, L. et al. Siderophores produced by Magnaporthe grisea in the presence and absence of iron. Z Naturforsch C 61, 461–464 (2006).

56 R., S. K. Studies on the physiology, morphology and serology of Exobasidium. Symb. Bot. Upsal. 18, 1–89 (1964).

57 Shannon, P. et al. Cytoscape: a software environment for integrated models of biomolecular interaction networks. Genome Res 11, 2498–2504 (2003).

