## Supplemental Information for "Native Electrospray-based Metabolomics Enables the Detection of Metal-binding Compounds"

### Supplementary Figures

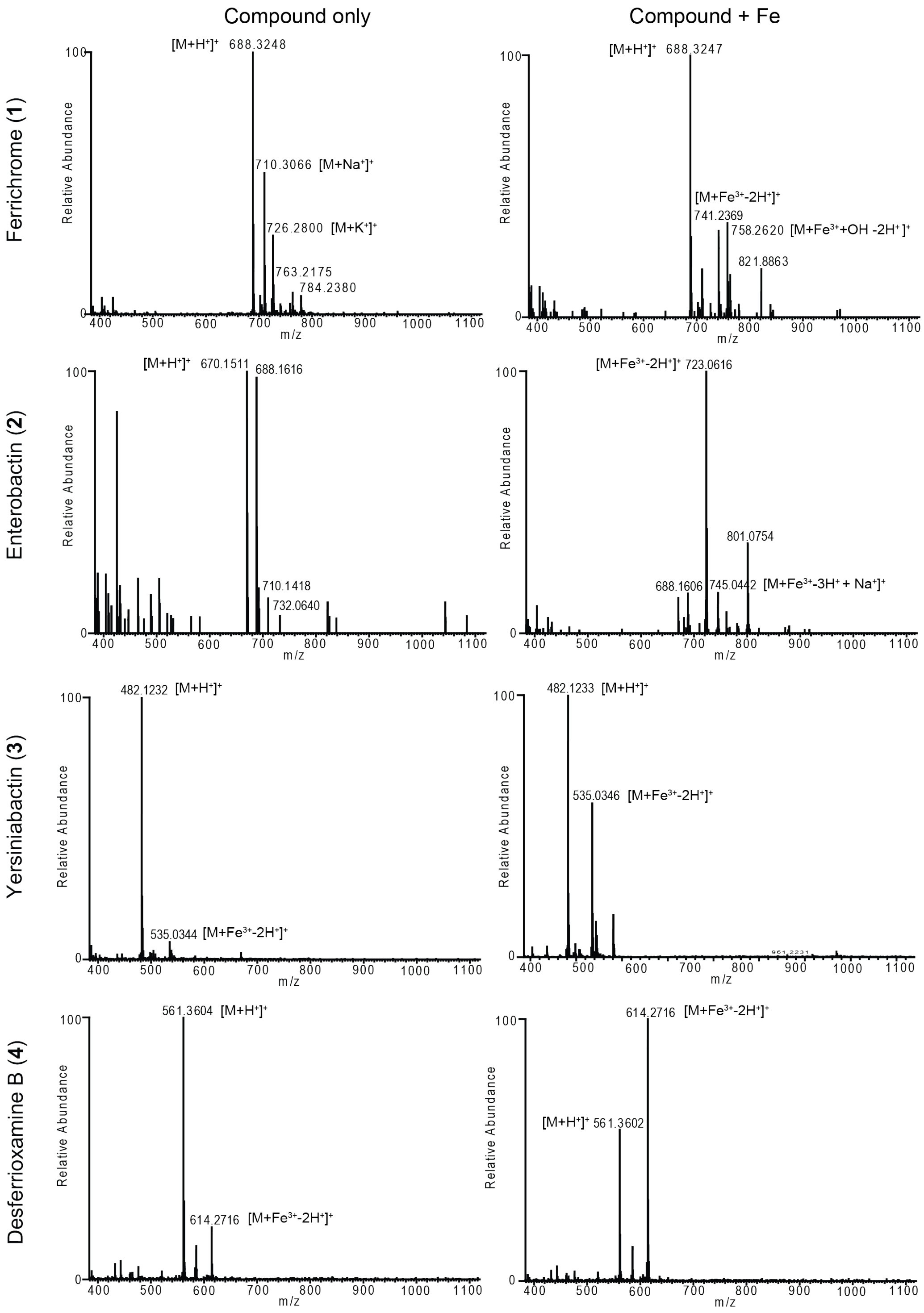


**SI Figure 1.** Iron-adducts are observed when iron salt is added to commercially available siderophores in 10 mM ammonium hydroxide at pH 6.8. Apo and iron-adducts are labeled for each compound.

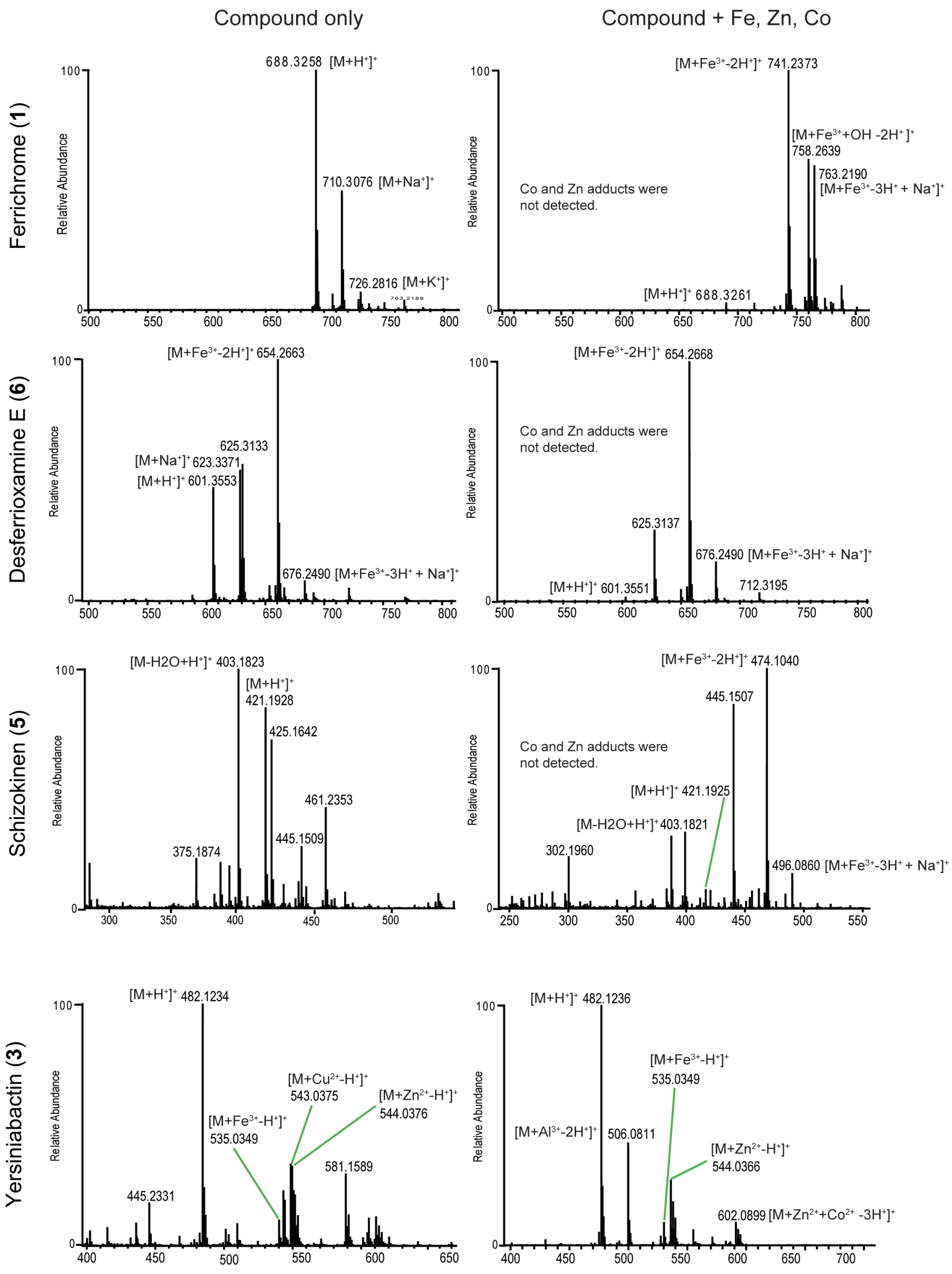


**SI Figure 2.** A number of adducts are observed when a combination of iron, cobalt, and zinc salts are added to commercially available siderophores in 10 mM ammonium hydroxide at pH 6.8. Apo and metal-bound adducts are labeled for each compound.

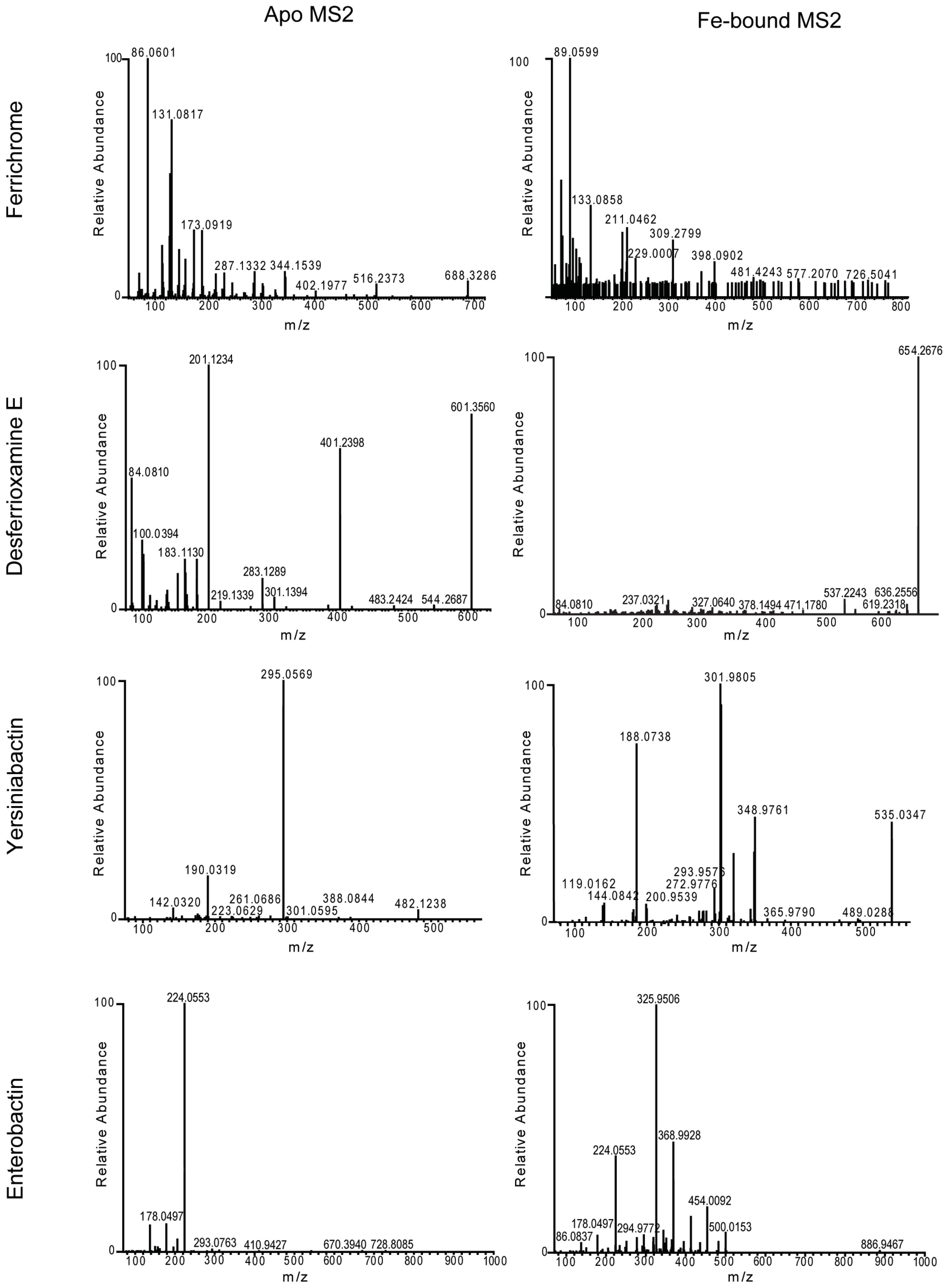


**SI Figure 3**. The $MS^2$ spectra of apo- ($[M+H^+]^+$) and metal-bound ($[M+Fe^{3+}-2H^+]^+$) siderophores differ significantly. As such, apo- and metal-bound species are often not connected by MS/MS similarity edges during molecular networking.

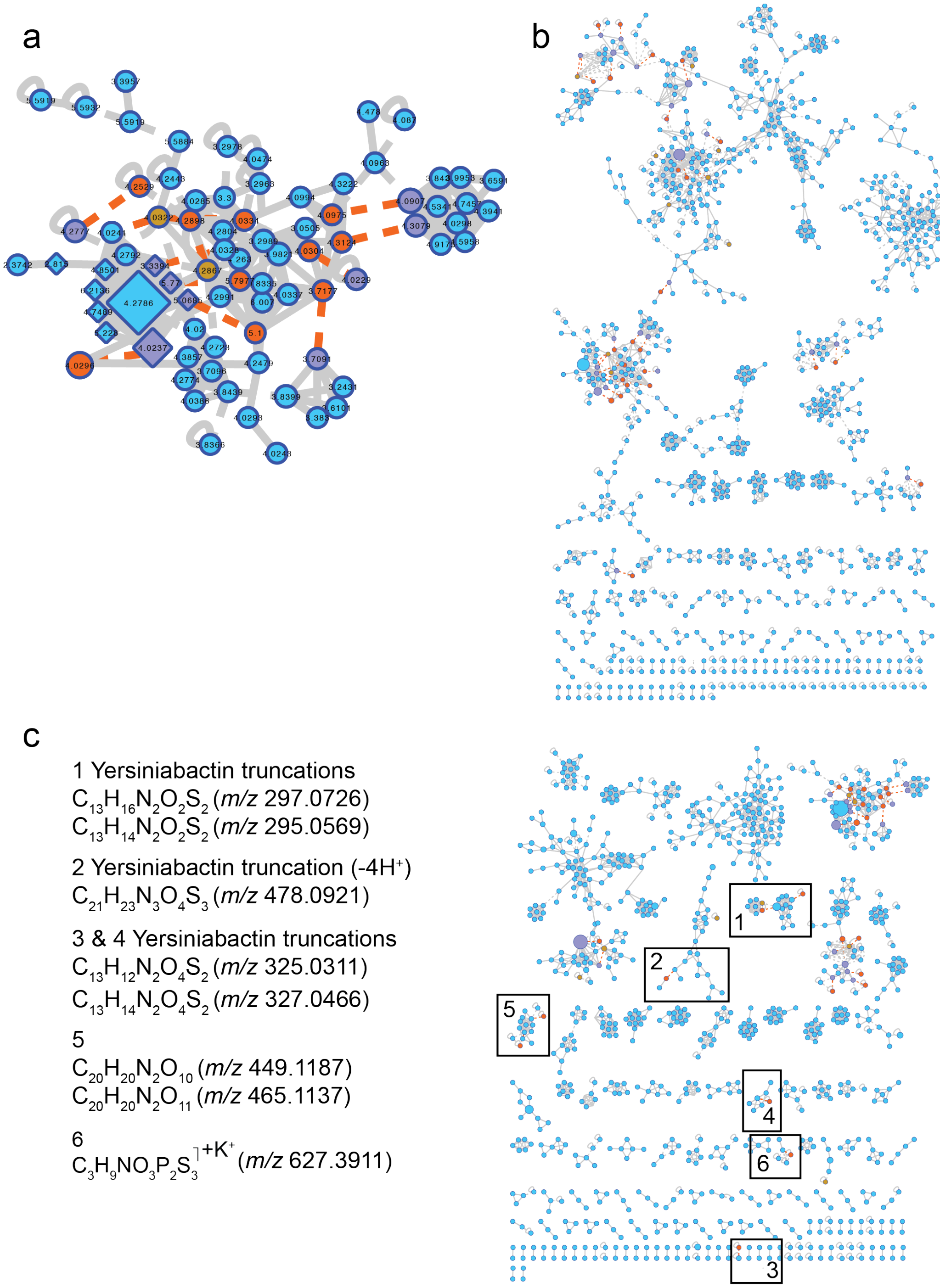


**SI Figure 4.** (a) The yersiniabactin molecular family highlighted in 3f is labeled with retention time rather than with precursor *m/z*. Fig 3f contains multiple nodes with the same precursor *m/z*, yet a number of these nodes elute at different retention times, suggesting multiple isomers are present. (b) Iron-binding molecules that do not cluster with the three molecular families highlighted in Fig 3f-h cluster together when IIN search includes additional adducts and in-source fragments. Additional adducts and in-source fragments searched for include M-H2O, M-2H2O, M-NH3, M-CO2, M+ACN, M+MeOH. (c) Iron-binding molecules that are present in molecular families other than those highlighted in Fig 3f-h have been assigned molecular formulas using SIRIUS 4.0[1]. The molecular formula for each precursor *m/z* was assigned based on the top score in SIRIUS 4.0[1].

### Supplementary Methods

**Iron addition to siderophore standards (direct infusion MS).** Yersiniabactin, ferrichrome, enterobactin, desferrioxamine B and E, and schizokinen (EMC Microcollections, https://www.microcollections.de/product-services/siderophores.html) were each dissolved to a 1 mM stock concentration in methanol (in the case of yersiniabactin, enterobactin, ferrioxamines, vibriobactin) or water (ferrichrome and rhodotorulic acid). Then 2 µL of this stock solution was added to each well in a 96-well plate for a 20 µM final stock. 97 µL of 10 mM ammonium acetate buffer was added to each well. "No metal" wells were prepared by adding an additional 1 µL of buffer, and "metal containing" wells were prepared by adding 1 µL of 20 mM metal solution stock (prepared in water) was added to yield a final concentration of 200 µM (for a 10-fold excess of metal). Mass spectrometry was performed as specified in the **Online Methods**; specifically, injections were performed in 95% A (10 mM ammonium acetate) at a flow rate of 0.40 mL/min.

**Mixed metal addition to siderophore standards (direct infusion MS).** Yersiniabactin, ferrichrome, desferrioxamine E (DFE), and schizokinen (EMC Microcollections, https://www.microcollections.de/product-services/siderophores.html) were each dissolved to a 1 mM stock concentration in methanol (in the case of yersiniabactin and desferrioxamine E) or water (ferrichrome and schizokinen). Then 0.5 µL of this stock solution was added to each well in a 96-well plate for a 10 µM final stock. 48.5 µL of 10 mM ammonium acetate buffer was added to each well. "No metal" wells were prepared by adding an additional 1 µL of buffer, and "metal-containing" wells were prepared to a final concentration of 2 µM for each metal or 8 µM for each metal. Metals were prepared as a 1 mM solution stock in water. Mass spectrometry was performed as specified in the **Online Methods**; specifically, injections were performed in 95% A (10 mM ammonium acetate) at a flow rate of 0.40 mL/min.

**Post LC-MS/MS pH neutralization and metal infusion set-up.** A 160 mM solution of $FeCl_3$ was prepared as the stock solution. Ammonium hydroxide solution at 1 M was prepared as the stock solution. Iron stock was diluted to final concentrations of 3.2 mM and 4.8 mM. Then siderophore mixture sample was run through a C18 column at a flow rate of 0.5 mL/min. Iron solutions of 3.2 and 4.8 mM were tested with a neutralizing solution of 1 M ammonium hydroxide at a flow rate of 5 µL/min. Post-LC pH was verified by collecting the flow through and spotting on pH paper (Sigma).

**Preparation of siderophore standard mixture (mixture A).** Sulfamethazine, sulfamethizole, sulfachloropyridazine, and sulfadimethoxine were prepared to a final concentration of 10 mg/L in a 1:1 water/methanol mixture (v/v), coumarin-314 was prepared to a final concentration of 20 mg/L in methanol, and amitryptiline was prepared to a final concentration of 10 mg/L in water. Ferrichrome was added from a 1 mM stock in water to a final concentration of 60 µM.

**Data analysis of ferrichrome in the presence of non-binding molecules (mixture A) - iron addition.** MS was run as described in LC-MS/MS data acquisition section. All raw and processed data is publicly available at ftp://massive.ucsd.edu/MSV000084237. MS/MS spectra were converted to .mzML files using MSconvert (ProteoWizard)[2]. MS1 feature extraction and MS/MS pairing was performed with the MZmine 2.37corr17.7_kai_merge2[3-5]. An intensity threshold of 5E5 for MS1 spectra and of 1E3 for MS/MS spectra was used. MS1 chromatogram building was performed within 10 ppm mass windows and a minimum peak intensity of 5E5 was set. Extracted Ion Chromatograms (XICs) were deconvoluted using the local minimum search algorithm with a chromatographic threshold of 0.01%, a search minimum in RT range of 0.2 min, and a median *m/z* center calculation with *m/z* range for MS2 pairing of 0.01 and RT range for MS2 scan pairing of 0.1. Isotope peaks were grouped and features from different samples were aligned with 10 ppm mass tolerance and 0.1 min retention time tolerance. MS1 peak lists were joined using an *m/z* tolerance of 10 ppm and retention time tolerance of 0.3 min; alignment was performed by placing a weight of 90 on m/z and 10 on retention time. Correlation of co-eluting features was performed with the metaCorrelate module; retention time tolerance of 0.15, minimum height of 0, noise level of 0 were used. A correlation of 85 was set as the cut off for the min feature shape corr and feature height correlation was not used. The following adducts were searched: $[M + H^+]^+$, $[M + Na^+]^+$, $[M + K^+]^+$, $[M + Ca^{2+}]^{2+}$, $[M + Fe^{3+} - 2H^+]^+$, with an m/z tolerance of 10 ppm, a maximum charge of 2, and maximum molecules/cluster of 2.

Peak areas and feature correlation pairs were exported as .csv files and the corresponding consensus MS/MS spectra were exported as an .mgf file. For spectral networking and spectrum library matching, all files were uploaded to the feature-based molecular networking workflow on GNPS (gnps.ucsd.edu, Wang et al. Nature Biotech, Aaron et al. Nature Protocols). The GNPS job can be accessed through the following link https://gnps.ucsd.edu/ProteoSAFe/status.jsp?task=79d0f380b4814ff9a720836c5570036f. All .csv and .mgf files in addition to MZmine 2 project can be accessed at ftp://massive.ucsd.edu/MSV000084237.

Molecular networks were visualized with Cytoscape 3.7.1[6]. Node size was scaled to feature abundances generated in MZmine 2[3-5], where node size was equal to summed feature abundance.

**Preparation of siderophore standard mixture (mixture B).** Five siderophores (ferrichrome, vibriobactin, yersiniabactin, enterobactin, and rhodotorulic acid) were dissolved in 50% MeOH/water. Siderophores were added at amounts that would result in similar peak heights - yersiniabactin was added to a 10 µM final concentration, ferrichrome was added to a 20 µM final concentration, and vibriobactin, rhodotorulic acid, and enterobactin were added to a 30 µM final concentration.

**Data analysis of standard mixture B - iron addition.** MS was run as described in LC-MS/MS data acquisition section. All raw and processed data is publicly available at ftp://massive.ucsd.edu/MSV000084237. MS/MS spectra were converted to .mzML files using MSconvert (ProteoWizard)[2]. MS1 feature extraction and MS/MS pairing was performed with the MZmine 2.37corr17.7_kai_merge2 [3-5]. An intensity threshold of 5E5 for MS1 spectra and of 1E3 for MS/MS spectra was used. MS1 chromatogram building was performed within 10 ppm mass windows and a minimum peak intensity of 5E5 was set. Extracted Ion Chromatograms (XICs) were deconvoluted using the local minimum search algorithm with a chromatographic threshold of 0.01%, a search minimum in RT range of 0.2 min, and a median *m/z* center calculation with *m/z* range for MS2 pairing of 0.01 and RT range for MS2 scan pairing of 0.1. Isotope peaks were grouped and features from different samples were aligned with 10 ppm mass tolerance and 0.1 min retention time tolerance. MS1 peak lists were joined using an *m/z* tolerance of 10 ppm and retention time tolerance of 0.1 min; alignment was performed by placing a weight of 90 on m/z and 10 on retention time. Gap filling was performed using an intensity tolerance of 10%, an m/z tolerance of 10 ppm, and a retention tolerance of 0.1. Correlation of co-eluting features was performed with the metaCorrelate module; retention time tolerance of 0.1, minimum height of 0, noise level of 0 were used. A correlation of 85 was set as the cut off for the min feature shape corr and feature height correlation was not used. The following adducts were searched: $[M + H^+]^+$, $[M + Na^+]^+$, $[M + K^+]^+$, $[M + Ca^{2+}]^{2+}$, $[M + Fe^{3+} - 2H^+]^+$, with an m/z tolerance of 10 ppm, a maximum charge of 2, and maximum molecules/cluster of 2.

Peak areas and feature correlation pairs were exported as .csv files and the corresponding consensus MS/MS spectra were exported as an .mgf file. For spectral networking and spectrum library matching, all files were uploaded to the feature-based molecular networking workflow on GNPS (gnps.ucsd.edu, Wang et al. Nature Biotech, Aaron et al. Nature Protocols). The GNPS job can be accessed through the following link https://gnps.ucsd.edu/ProteoSAFe/status.jsp?task=ad4b2665dfb744d09a9d2445f1213720. All .csv and .mgf files in addition to the MZmine 2 project can be accessed at ftp://massive.ucsd.edu/MSV000084237.

Molecular networks were visualized with Cytoscape 3.7.1[6]. Node size was scaled to feature abundances generated in MZmine 2, where node size was equal to feature abundance in post-LC Fe-infusion run.

**Preparation of siderophore standard mixture (mixture C).** Four siderophores (ferrichrome, vibriobactin, yersiniabactin, and rhodotorulic acid) were dissolved in 50% MeOH/water. Siderophores were added at amounts that would result in similar peak heights - yersiniabactin was added to a 10 μM final concentration, ferrichrome was added to a 20 μM final concentration, and vibriobactin and rhodotorulic acid were added to a 30 μM final concentration.

**Data analysis of standard mixture C - iron, cobalt, and zinc addition.** MS was run as described in LC-MS/MS data acquisition section. All raw and processed data is publicly available at ftp://massive.ucsd.edu/MSV000084237. MS/MS spectra were converted to .mzML files using MSconvert (ProteoWizard)[2]. MS1 feature extraction and MS/MS pairing was performed with the MZmine 2.37corr17.7_kai_merge2[3-5]. An intensity threshold of 3E5 for MS1 spectra and of 1E3 for MS/MS spectra was used. Crop filter was applied to exclude features after 6 minutes. MS1 chromatogram building was performed within a 10 ppm mass windows and a minimum peak intensity of 3E5 was set. Extracted Ion Chromatograms (XICs) were deconvoluted using the local minimum search algorithm with a chromatographic threshold of 0.01%, a search minimum in RT range of 0.2 min, and a median m/z center calculation with *m/z* range for MS2 pairing of 0.01 and RT range for MS2 scan pairing of 0.1. Isotope peaks were grouped and features from different samples were aligned with 10 ppm mass tolerance and 0.1 min retention time tolerance. MS1 peak lists were joined using an m/z tolerance of 10 ppm and retention time tolerance of 0.1 min; alignment was performed by placing a weight of 90 on *m/z* and 10 on retention time. Gap filling was performed using an intensity tolerance of 10%, an *m/z* tolerance of 10 ppm, and a retention tolerance of 0.1. Correlation of co-eluting features was performed with the metaCorrelate module; retention time tolerance of 0.1, minimum height of 0, noise level of 0 were used. A correlation of 85 was set as the cut off for the min feature shape corr and feature height correlation was not used. The following adducts were searched: $[M + H^+]^+$, $[M + Na^+]^+$, $[M + K^+]^+$, $[M + Ca^{2+}]^{2+}$, $[M + Fe^{3+} - 2H^+]^+$, $[M + Zn^{2+} - H^+]^+$, $[M + Co^{2+} - H^+]^+$ with an *m/z* tolerance of 10 ppm, a maximum charge of 2, and maximum molecules/cluster of 2.

Peak areas and feature correlation pairs were exported as .csv files and the corresponding consensus MS/MS spectra were exported as an .mgf. file. For spectral networking and spectrum library matching, all files were uploaded to the feature-based molecular networking workflow on GNPS (gnps.ucsd.edu, Wang et al. Nature Biotech, Aaron et al. Nature Protocols). The GNPS job can be accessed through the following link http://gnps.ucsd.edu/ProteoSAFe/status.jsp?task=0d8cacfd74744357ada2f7a081252d64. All .csv and .mgf files in addition to MZmine 2 project can be accessed at ftp://massive.ucsd.edu/MSV000084237.

Molecular networks were visualized with Cytoscape 3.7.1[6]. Node size was scaled to feature abundances generated in MZmine 2, where node size was equal to feature abundance in post-LC Fe-infusion run.

**Preparation of cheese culture extracts.** Extracts were prepared from *Glutamicibacter arilaitensis JB182*[7] grown in liquid cheese media. All glassware was washed with 6 molar (M) aqueous HCl and rinsed with deionized water. *Glutamicibacter airlaitensis JB182* was grown from glycerol stocks on lysogeny broth agar at room temperature for two days. 100 mL of liquid cheese curd medium (2% (w/v) freeze-dried Bayley cheese curd (Jasper Hill Creamery, Greensboro Bend, VT), 3% (w/v) NaCl, pH 7) were inoculated from single colonies of JB182. Cultures were grown in triplicate at room temperature for seven days shaking at 125 revolutions per minute (rpm). 2 x 50 ml of each culture were transferred to 50 ml conical tubes and centrifuged in an Eppendorf 5810 R centrifuge (Eppendorf, Hamburg, Germany) at 3900 rpm for 45 minutes. Supernatants were filter-sterilized and stored at -80 ºC. Thawed supernatants were applied to HypersepTMC18 cartridges (Thermo Fisher Scientific, Waltham, MA) that were preconditioned with 12 ml of 100% (volume/volume) methanol (HPLC grade, 99.9% purity, Fisher Scientific, Hampton, NH) followed by 12 ml of UltraPure distilled water (Thermo Fisher Scientific, Waltham, MA). Columns were then washed with 6 ml of 5% (v/v) aqueous methanol and eluted with 1 ml of 40% (v/v) aqueous methanol, followed by 1 ml of 60% (v/v) aqueous methanol, 1 ml of 80% (v/v) aqueous methanol, and lastly 1 ml of 100% methanol. Each fraction was collected separately and stored at -20ºC.

**Data analysis of cheese culture extracts - iron addition.** MS was run as described in LC-MS/MS data acquisition section. All raw and processed data is publicly available at ftp://massive.ucsd.edu/MSV000084289/. MS/MS spectra were converted to .mzML files using MSconvert (ProteoWizard)[2]. MS1 feature extraction and MS/MS pairing was performed with the MZmine 2.37corr17.7_kai_merge2[3-5]. An intensity threshold of 1E5 for MS1 spectra and of 1E3 for MS/MS spectra was used. MS1 chromatogram building was performed within a 10 ppm mass windows and a minimum peak intensity of 1E5 was set. Extracted Ion Chromatograms (XICs) were deconvoluted using the local minimum search algorithm with a chromatographic threshold of 0.01%, a search minimum in RT range of 0.2 min, and a median m/z center calculation with m/z range for MS2 pairing of 0.01 and RT range for MS2 scan pairing of 0.1. Isotope peaks were grouped and features from different samples were aligned with 10 ppm mass tolerance and 0.1 min retention time tolerance. MS1 peak lists were joined using an m/z tolerance of 10 ppm and retention time tolerance of 0.1 min; alignment was performed by placing a weight of 75 on m/z and 25 on retention time. Gap filling was performed using an intensity tolerance of 10%, an m/z tolerance of 10 ppm, and a retention tolerance of 0.1. Correlation of co-eluting features was performed with the metaCorrelate module; retention time tolerance of 0.1, minimum height of 1E5, noise level of 1E4 were used. A correlation of 85 was set as the cut off for the min feature shape correlation and feature height correlation was not used. The following adducts were searched: $[M + H^+]^+$, $[M + Na^+]^+$, $[M + K^+]^+$, $[M + Ca^{2+}]^{2+}$, $[M + Fe^{3+} - 2H^+]^+$, with an m/z tolerance of 10 ppm, a maximum charge of 2, and maximum molecules/cluster of 2.

Peak areas and feature correlation pairs were exported as .csv files and the corresponding consensus MS/MS spectra were exported as an .mgf. file. For spectral networking and spectrum library matching, all files were uploaded to the feature-based molecular networking workflow on GNPS (gnps.ucsd.edu, Wang et al. Nature Biotech, Aaron et al. Nature Protocols). For spectrum library matching and spectral networking the minimum cosine score to define spectral similarity was set to 0.7. The Precursor and Fragment Ion Mass Tolerances were set to 0.01 Da and Minimum Matched Fragment Ions to 4, Minimum Cluster Size to 1 (MS Cluster off). When Analog Search was performed the maximum mass difference was set to 100 Da. The GNPS job for the cheese culture can be accessed through the following link https://gnps.ucsd.edu/ProteoSAFe/status.jsp?task=5459d22126e843a3a1449f8362cd267f. All .csv and .mgf files in addition to the MZmine 2 project can be accessed at ftp://massive.ucsd.edu/MSV000084289.

Molecular networks were visualized with Cytoscape 3.7.1[6]. Node size was scaled to the summed precursor intensity, where node size was scaled based on feature abundance summed across the runs.

**Preparation of *E. coli* Nissle culture extracts.** Wild-type *E. coli* Nissle 1917 was pre-cultured in 5ml LB broth overnight in triplicate at 37 °C with agitation, followed by two-times of wash with M9 glucose minimal medium (prepared by adding the following per 1 L: 7.5g $Na_2HPO_4$, 3g $KH_2PO_4$, 0.5g NaCl, 1g $NH_4Cl$, 0.1mM CaCl2, 0.5mM $MgSO_4$, 0.2% glucose). Cell pellets were resuspended in M9 and inoculated into 20ml M9 medium for overnight growth at 37 °C with agitation. Cells were then centrifuged down and washed twice in M9, and inoculated into 30ml M9 for overnight growth at 37 °C with agitation. Then the cells were spun down, and supernatant was collected for analysis.

**Data analysis of *E. coli* Nissle extracts – iron addition.** MS was run as described in LC-MS/MS data acquisition section. MS/MS spectra were converted to .mzML files using MSconvert (ProteoWizard)[2]. All raw and processed data is publicly available at ftp://massive.ucsd.edu/MSV000083387/. MS1 feature extraction and MS/MS pairing was performed with MZmine 2.37corr17.7_kai_merge2[3-5]. An intensity threshold of 1E6 for MS1 spectra and of 1E3 for MS/MS spectra was used. MS1 chromatogram building was performed within a 10 ppm mass windows and a minimum peak intensity of 3E5 was set. Extracted Ion Chromatograms (XICs) were deconvoluted using baseline cutoff with m/z range for MS2 pairing of 0.01 and RT range for MS2 scan pairing of 0.2. After chromatographic deconvolution, MS1 features linked to MS/MS spectra within 0.01 m/z mass and 0.2 min retention time windows. Isotope peaks were grouped and features from different samples were aligned with 10 ppm mass tolerance and 0.1 min retention time tolerance. MS1 peak lists were joined using an m/z tolerance of 10 ppm and retention time tolerance of 0.1 min; alignment was performed by placing a weight of 75 on m/z and 25 on retention time. Gap filling was performed using an intensity tolerance of 10%, an m/z tolerance of 10 ppm, and a retention tolerance of 0.1. Correlation of co-eluting features was performed with the metaCorrelate module; retention time tolerance of 0.1, minimum height of 1E5, noise level of 1E4 were used. A correlation of 85 was set as the cut off for the min feature shape corr., and feature height correlation was not used. The following adducts were searched: $[M + H^+]^+$, $[M + Na^+]^+$, $[M + K^+]^+$, $[M + Ca^{2+}]^{2+}$, $[M + Fe^{3+} - 2H^+]^+$, with an m/z tolerance of 10 ppm, a maximum charge of 2, and maximum molecules/cluster of 2.

Peak areas and feature correlation pairs were exported as .csv files and the corresponding consensus MS/MS spectra were exported as an .mgf. file. For spectral networking and spectrum library matching, all files were uploaded to the feature-based molecular networking workflow on GNPS (gnps.ucsd.edu)[8,9]. For spectrum library matching and spectral networking the minimum cosine score to define spectral similarity was set to 0.7. The Precursor and Fragment Ion Mass Tolerances were set to 0.01 Da and Minimum Matched Fragment Ions to 4, Minimum Cluster Size to 1 (MS Cluster off). When Analog Search was performed the maximum mass difference was set to 100 Da. The GNPS job for the siderophore mix can be accessed: http://gnps.ucsd.edu/ProteoSAFe/status.jsp?task=1c3e79f0ab984386bd468e2d163281e0. All .csv and .mgf files in addition to the MZmine 2 project can be accessed at ftp://massive.ucsd.edu/MSV000083387. Mgf files were exported for SIRIUS in MZmine2, then molecular formulas were determined using SIRIUS 4.0.1 (build 9)[1].

Molecular networks were visualized with Cytoscape 3.7.1[6]. Node size was scaled to the summed precursor intensity, where node size was scaled based on feature abundance summed across the runs.

**Preparation of *Eutypa lata* extracts.** *Eutypa lata* was grown in liquid YMG medium (4 g/L yeast extract, 10 g/L malt extract, 10 g/L glucose) for five days. Mycelium was washed three times with Sundström minimal medium[10] and transferred to Erlenmeyer flasks with Sundström minimal medium. All liquid cultivations were carried out with agitation at 120 rpm on an orbital shaker at room temperature[11]. After 18 days cultivation in Sundström medium, XAD-16 (10% w/v) were added to culture filtrate and incubated for two hours with agitation at 120 rpm. The resin was removed by filtration and washed with three volumes of water. Then metabolites were eluted by methanol. The eluate was concentrated by evaporation to dryness and stored at -80 °C for analysis.

**Data analysis of *Eutypa lata* extracts – iron extracts.** MS was run as described in LC-MS/MS data acquisition section. MS/MS spectra were converted to .mzML files using MSconvert (ProteoWizard)[2]. All raw and processed data is publicly available at ftp://massive.ucsd.edu/MSV000084030/. MS1 feature extraction and MS/MS pairing was performed with MZmine 2.37corr17.7_kai_merge2[3,4]An intensity threshold of 1E6 for MS1 spectra and of 1E3 for MS/MS spectra was used. MS1 chromatogram building was performed within a 10 ppm mass windows and a minimum peak intensity of 3E5 was set. Extracted Ion Chromatograms (XICs) were deconvoluted using the local minimum search algorithm with a chromatographic threshold of 0.01%, a search minimum in RT range of 0.1 min, and a median m/z center calculation with m/z range for MS2 pairing of 0.01 and RT range for MS2 scan pairing of 0.2. After chromatographic deconvolution, MS1 features linked to MS/MS spectra within 0.01 m/z mass and 0.2 min retention time windows. Isotope peaks were grouped and features from different samples were aligned with 10 ppm mass tolerance and 0.1 min retention time tolerance. MS1 peak lists were joined using an m/z tolerance of 10 ppm and retention time tolerance of 0.1 min; alignment was performed by placing a weight of 75 on m/z and 25 on retention time. Gap filling was performed using an intensity tolerance of 10%, an m/z tolerance of 10 ppm, and a retention tolerance of 0.1. Correlation of co-eluting features was performed with the metaCorrelate module; retention time tolerance of 0.1, minimum height of 1E5, noise level of 1E4 were used. A correlation of 0.85 was set as the cut off for the min feature shape corr. The following adducts $[M + H^+]^+$, $[M + Na^+]^+$, $[M + K^+]^+$, $[M + Ca^{2+}]^{2+}$, $[M + Fe^{3+} - 2H^+]^+$, in combination with the neutral loss of water for each adduct [M-H2O+X] (where X is one of the listed adducts) were searched with an m/z tolerance of 10 ppm, a maximum charge of 2, and maximum molecules/cluster of 2.

Peak areas and feature correlation pairs were exported as .csv files and the corresponding consensus MS/MS spectra were exported as an .mgf. file. For spectral networking and spectrum library matching, all files were uploaded to the feature-based molecular networking workflow on GNPS (gnps.ucsd.edu)[8,9]. For spectrum library matching and spectral networking, the minimum cosine score to define spectral similarity was set to 0.7. The Precursor and Fragment Ion Mass Tolerances were set to 0.01 Da and Minimum Matched Fragment Ions to 4, Minimum Cluster Size to 1 (MS Cluster off). When Analog Search was performed the maximum mass difference was set to 100 Da. The GNPS job for the wine fungus can be accessed http://gnps.ucsd.edu/ProteoSAFe/status.jsp?task=256ba734f4334c1c90f65ffbd9141d0e. All .csv and .mgf files in addition to the MZmine 2 project can be accessed at ftp://massive.ucsd.edu/MSV000084030/. Mgf files were exported for SIRIUS in MZmine2, then molecular formulas were determined using SIRIUS 4.0.1 (build 9)[1].

Molecular networks were visualized with Cytoscape 3.7.1[6]. Node size was scaled to the summed precursor intensity, where node size was scaled based on feature abundance summed across the runs.


1 Dührkop, K. *et al.* SIRIUS 4: a rapid tool for turning tandem mass spectra into metabolite structure information. *Nat Methods* **16**, 299-302 (2019).

2 Kessner, D., Chambers, M., Burke, R., Agus, D. & Mallick, P. ProteoWizard: open source software for rapid proteomics tools development. *Bioinformatics* **24**, 2534–2536 (2008).

3 Pluskal, T., Castillo, S., Villar-Briones, A. & Orešič, M. MZmine 2: Modular framework for processing, visualizing, and analyzing mass spectrometry-based molecular profile data. *BMC Bioinformaticsvolume* **11**, 395 (2010).

4 Katajamaa, M., Miettinen, J. & Oresic, M. MZmine: toolbox for processing and visualization of mass spectrometry based molecular profile data. *Bioinformatics* **22**, 634-636 (2).

5 Schmid, R. *et al.* Ion identity based molecular networking. *under revision* (2019).

6 Shannon, P. *et al.* Cytoscape: a software environment for integrated models of biomolecular interaction networks. *Genome Res* **11**, 2498-2504 (2003).

7 Bonham, K. S., Wolfe, B. E. & Dutton, R. J. Extensive horizontal gene transfer in cheese-associated bacteria. *eLife* **6**, e22144, doi:10.7554/eLife.22144 (2017).

8 Wang, M. Sharing and community curation of mass spectrometry data with Global Natural Products Social Molecular Networking. *Nat Biotechnol.* **34**, 828-837 (2016).

9 Aron, A. T. *et al.* Reproducible Molecular Networking Of Untargeted Mass Spectrometry Data Using GNPS. *ChemRxiv Preprint*, doi:10.26434/chemrxiv.9333212.v1 (2019 ChemRxiv. Preprint).

10 R., S. K. Studies on the physiology, morphology and serology of Exobasidium. *Symb. Bot. Upsal.* **18**, 1-89 (1964).

11 Antelo, L. *et al.* Siderophores produced by Magnaporthe grisea in the presence and absence of iron. *Z Naturforsch C* **61**, 461-464 (2006).
